# Mutations in ASH1L cause a neurodevelopmental disorder with sex differences in epilepsy and autism

**DOI:** 10.1101/2025.02.21.639570

**Authors:** Carin M. Papendorp, Elizabeth Nolan, Haruki Higashimori, Megha Jhanji, Luis Goicouria, Cosmo A. Pieplow, Kirsten Whitley, Steven Defreitas, Jennifer Elacio, Duyu Nie, Brian Kavanaugh, Carrie R. Best, Sofia B. Lizarraga, Judy S. Liu

## Abstract

To understand brain phenotypes associated with ASH1L, we performed both studies in mouse models and clinical phenotyping of human subjects. We found in mice that ASH1L mutations result in seizures, microcephaly, and also less complex dendritic morphology. When we analyzed human subjects based for epilepsy, intellectual disability, and ASD, we found sex differences in epilepsy and autism, with epilepsy predominantly in female and ASD in male subjects. To understand the cellular and molecular mechanisms of the sex-difference, we performed whole cell patch clamp electrophysiology in mice and found hyperexcitability in female compared with male hippocampal CA1 neurons. We report the identification of sex-specific transcriptomic signatures resulting from ASH1L haploinsufficiency. Differentially expressed genes in female mice showed distinct association with epileptic encephalopathy, postnatal microcephaly and autistic behaviors. Thus, the role of ASH1L in specific circuits may be sex-dependent leading to sexual dimorphic effects from disruption of this gene.

## Introduction

In recent years, large sequencing efforts have identified highly penetrant variants in individual genes, including the gene *Absent, small, or homeotic-like* (*ASH1L*), as risk factors for neurodevelopmental disorders. These studies focus on specific disorders for example, autism spectrum disorder (ASD)^1,2^, epilepsy^3,4^, schizophrenia, or Tourette’s syndrome^5^. These large sequencing efforts are often not accompanied by sufficient clinical data, thus, the clinical phenotypes for these newly identified causative genes such as *ASH1L* have not been fully defined. For many of these newly identified disease genes, including *ASH1L*, there are currently just a handful of case reports in the literature^6–11^. However, given the increase in clinical sequencing, new patients are rapidly being diagnosed, and there exists an urgent need to perform both clinical and basic science studies.

Since the mammalian *ASH1L* gene was first cloned in 2000^12^, *in vitro* experiments have yielded insights regarding the basic molecular function of the ASH1L enzyme. ASH1L is a histone lysine methyltransferase^13^ thought to methylate the histone lysine residue on histone 3: lysine 36 (H3K36).^14–16^ Methylation at this lysine residues is essential for transcriptional initiation and elongation.^17^ ASH1L is ubiquitously expressed throughout the body.^18–21^ In the brain, ASH1L is expressed across brain areas^22^ and cell types, including excitatory and inhibitory neurons, astrocytes, oligodendrocytes, and microglia.^23–26^ and not preferentially expressed in any specific brain region. In humans, *ASH1L* mRNA expression levels are fairly equal across all regions of cortex. ^27,28^

ASH1L has been studied in rodent brain and other systems, however models that have been used include a heterozygous loss of function mutation^29^, conditional deletion in the brain ^29,30^, and shRNAi for acute knockdown^31^, which may not fully replicate the entire scope of mutations leading to the human condition. In patients, ASH1L mutations are heterozygous, where only one copy is affected. This is thought to reflect a genetic mechanism of haploinsufficiency ^32^. The types of mutations in *ASH1L* that have brain effects are diverse, including missense and truncations occurring throughout the gene. In order to understand to role of ASH1L in patients with deleterious genetic changes, we planned a human clinical phenotyping study. We also studied a rodent model of ASH1L across a standard inbred mouse strain, C57BL/6J(B6), and an outbred strain, CD-1. We then compared findings in our mouse models to the results of the first human clinical phenotyping study of ASH1L where we characterized twenty-three previously unreported patients with ASH1L. We find epilepsy more often in female subjects in our cohort, that has not been previously reported. We are also able demonstrate hyper-excitability in hippocampal CA1 neurons by whole cell-patch clamp electrophysiology specifically in female mice. The clinical presentation and electrophysiological studies in mice suggest there are distinct underlying mechanisms in males and females in response to ASH1L dysfunction.

We report the identification of sex-specific transcriptomic signatures that are genotype dependent with respect to the regulation of myelination, autophagy, and protein folding. Differentially expressed genes in heterozygous male and female mice showed distinct association with epileptic encephalopathy, postnatal microcephaly and autistic behaviors. Weighted gene co-expression analysis (WGCNA) defined additional molecular signatures in males and females distinctly modulated by ASH1L that are essential during development of neuronal circuits, that might be cell type specific and that could underlie the genetic risk of complex brain disorders such as autism and schizophrenia.

## Results

### Genetic background influences survival in ASH1L mutant mice

For this study, we used mice harboring a gene-trap (gt) mutation in *Ash1l*. A premature poly-A tail is inserted after the first translated exon, rendering the mice hypomorphic for ASH1L expression (Figure 1A).^33,34^ The homozygous *Ash1l^gt/gt^* mouse has a ∼90% decrease in *Ash1l* mRNA by reverse transcriptase quantitative polymerase chain reaction.^33,34^ Similarly, our data confirm a decrease in (though not total elimination of) *Ash1l* mRNA expression in the cortex and hippocampus in the homozygous *Ash1l^gt/gt^* mouse (Figure 1B).

**Figure 1:**
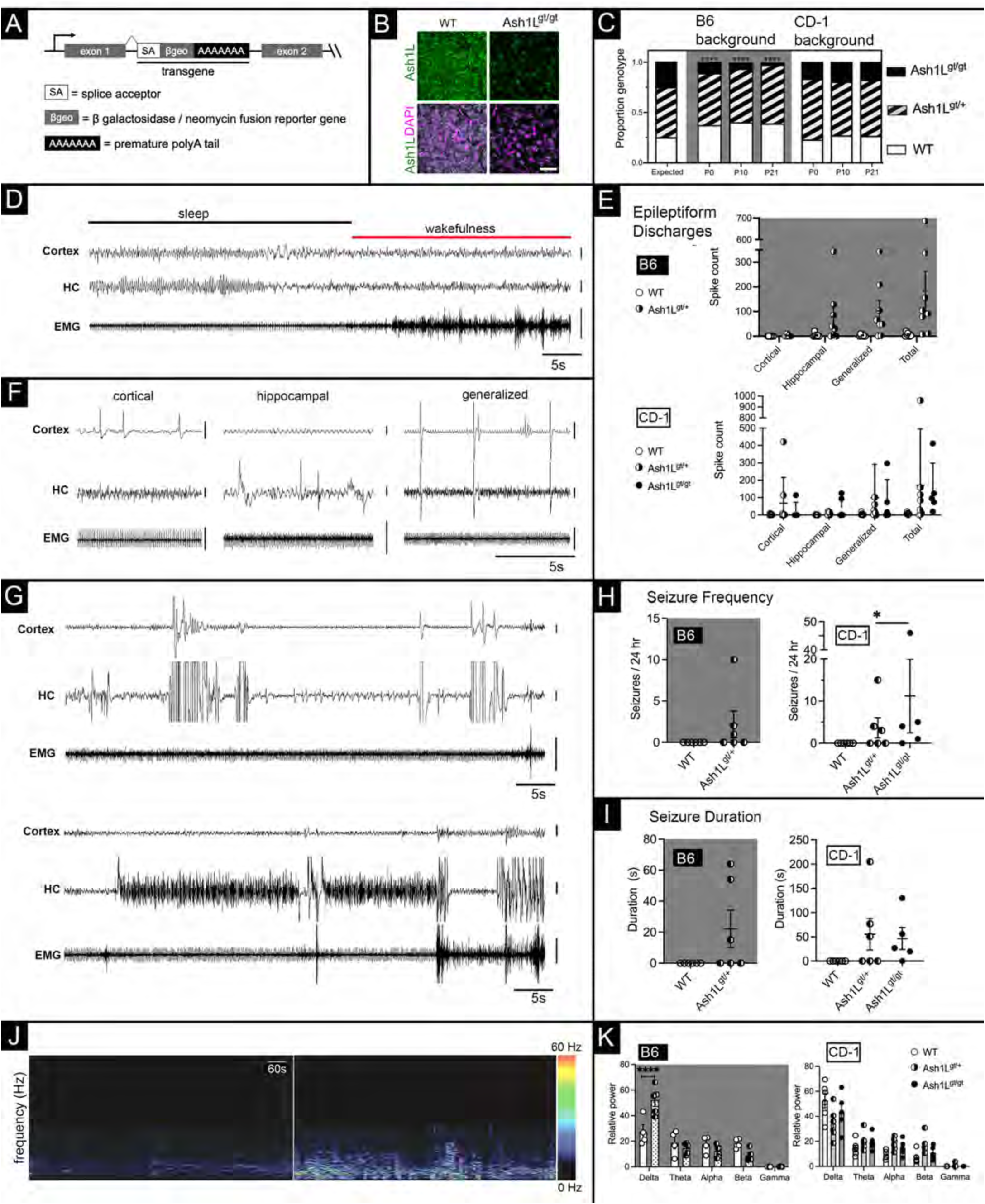
ASH1L mutant mice have epileptiform discharges, seizures, and altered power spectrum by EEG. (A) Schematic showing the gene-trap transgene inserted after the first translated exon of the ASH1L gene. The transgene consists of a splice acceptor (SA), β-galactosidase reporter, and a premature poly-A tail. (B) In-situ hybridization for ASH1L (green) on brain sections shows that *Ash1L* mRNA expression is reduced, but not absent, in the *Ash1L^gt/gt^* cortex. The counterstain is DAPI (bottom panels) in magenta. Scale bar = 50 μm. (C) Genetic background affects the Mendelian ratios of ASH1L mutant mice. Founder *Ash1L^gt/+^* mice were backcrossed more than three times to WT B6 and CD-1 mice to generate experimental animals. Experimental animals were analyzed at three time points: postnatal day zero (P0), postnatal day 14 (P14), and 2 months (2M). On a B6 background, *Ash1L^gt/gt^* mice are not born at the expected Mendelian ratios. *Ash1L^gt/gt^* that are born die around weaning age. Data were analyzed by Chi-square test at P0 (22.74, 2), P10 (25.50, 2), and P21 (34.61, 2). Expected Mendelian ratios (25% WT, 50% *Ash1L^gt/+,^* 25% *Ash1L^gt/gt^*) are shown on the left. On a CD-1 background, there is no significant difference between the expected and observed Mendelian ratios. Data were analyzed by Chi-square test at P0 (3.352, 2), P10 (2.556, 2), and P21 (3.292, 2). *P<.05, **P<.01, ***P<.001, ****P<.0001. (D) In a wild type B6 mouse, a normal transition between sleep and wakeful state is shown. Electrodes are placed in the L frontal cortex (top), R hippocampus (HC, middle), and trapezius muscle to record an electromyograph (EMG, bottom). Vertical scale bars (D, F, G) are 100μV and horizontal bars are 5 seconds. (E-F) Interictal epileptiform discharges or spikes were found to arise from the cortex, hippocampus, and also in a generalized manner. On both the B6 (E) and CD-1 (F) background, ASH1L mutant mice have interictal spikes. On the B6 background, the spikes are predominantly hippocampal and generalized, while some CD-1 mice have cortical spiking in addition to generalized and hippocampal epileptiform discharges. (G) Two examples of focal hippocampal seizures during sleep in a B6 (top) and CD-1 (bottom) mouse. There is baseline sleep activity in the cortical electrode (cortex) and electromyograph (EMG, black). The hippocampal electrode (HC) shows seizure activity. On a B6 background, approximately half of two-month-old *Ash1L^gt/+^*mice have electroencephalographic seizures. Seizure frequency (H) and average seizure duration (I) are calculated over a 24-hour period. No WT C57BL/6 mice have seizures. N = 6 mice per genotype (3 males, 3 females). (E-F) On a CD-1 background, approximately half of two-month-old *Ash1L^gt/+^* mice and most *Ash1L^gt/gt^* mice have electroencephalographic seizures. (J) Representative spectrograms of the cortical electrode from a WT (left) and *Ash1L^gt/+^* (right) mouse showing EEG frequency (Hz) vs. power (μV^2^) over 15 minutes of recording. The *Ash1L^gt/+^* mouse shows increased power in low frequencies. (K) On a B6 background, *Ash1L^gt/+^* mice have a significant increase in relative power in the delta frequency band. Delta = 0.5-4 Hz, Theta = 5.5-8.5 Hz, Alpha = 8-13 Hz, Beta= 13-30 Hz, Gamma = 35-44 Hz. N = 4 mice per genotype. On a CD-1 background, ASH1L mutant mice have no significant differences in relative EEG power. N = 6 mice per genotype.

In order to study the effect of ASH1L across genotypes, we backcrossed mice with the *Ash1l^g^*^t^ allele into either C57BL/6J (B6) background or to the outbred CD-1 strain. On the B6 background, homozygous loss of *Ash1l* is lethal perinatally ^5,14,33^ indicating that loss of ASH1L severely disrupts development (Figure 1C left). However, while some have reported that homozygous *Ash1l* mutations are embryonic lethal, on an inbred C57BL/6J (B6) background ^14^, we observe a combination of embryonic and postnatal mortality (Figure 1C). Homozygous *Ash1l^gt/gt^* mice are not born at the expected Mendelian ratios, but some *Ash1l^gt/gt^* survive through the early postnatal period (Figure 1C). However, all surviving *Ash1l^gt/gt^* mice die prior to weaning or shortly thereafter.

Surprisingly, when backcrossed for three generations to the outbred CD-1 strain, *Ash1l^gt/gt^*mice survive to adulthood and are fertile. On a CD-1 background, offspring from a *Ash1l^gt/+^* x *Ash1l^gt/+^* cross are born at the expected Mendelian ratios (Figure 1C, right). Additionally, while crossing founder hybrid mice to a B6 background decreases the proportion of *Ash1l^gt/gt^* mice, crossing to a CD-1 background increases the proportion of *Ash1l^gt/gt^* mice (Figure 1C). Clearly, genetic background exerts a profound influence on phenotype in the ASH1L mutant mouse. Therefore, we chose to examine phenotypes on both the inbred B6 background and the outbred CD-1 background.

### ASH1L mice have seizures, interictal discharges and altered power spectra by EEG

We performed video EEG monitoring in *Ash1l^gt/+^* mice on both backgrounds and found that disruption of ASH1L causes seizures. Wild type mice lack seizures or many epileptiform discharges on the B6 background. Based on EEG and EMG traces, sleep and wake states are easily appreciated (Figure 1D). On both backgrounds, EEG recordings detect interictal epileptiform discharges (spikes) on both backgrounds when ASH1L is disrupted (Figure E, F). We quantified only those EEG changes that were clearly not a result of artifact or movement, ie. single spikes with an amplitude of 2x baseline in the absence of EMG activity and without visible normal movement by the mouse (e.g. scratching, chewing, walking) associated with the EEG change. In the BL6 background, epileptiform discharges were detected mostly in leads implanted in the hippocampus, or simultaneously in both contralateral hippocampal and cortical leads ie, they were generalized (Figure 1E top and F). CD-1 mice had cortical spikes while cortical spikes were basically absent in B6 mice (Figure 1E).

While WT mice have no seizures, one-half of B6 *Ash1l^gt/+^* mice, one-half of CD-1 *Ash1l^gt/+^*mice, and 4 out of 5 CD-1 *Ash1l^gt/gt^* mice have electroencephalographic seizures (Figure 1G, H, I). Both male and female mutant mice have seizures. Seizures arise predominantly from the hippocampus during sleep, with sudden high-amplitude activity without activation in the contralateral cortical lead and without a behavioral correlate (Figure 1G). However, some generalized seizures are also observed. One CD-1 *Ash1l^gt/gt^*mouse appeared to die directly after a seizure. In these mice, the average seizure duration is less than one minute (Figure 1I). While mice occasionally have convulsive seizures, most electroencephalographic seizures are associated with freezing or behavioral pause (Supplemental Movies 2 and 3).

Changes in brain circuits manifest as differences in the power spectrum density of the EEG signal^35^. Alterations in spectral density are common in ASD and Neurodevelopmental disorders (NDD) and may be an important biomarker of ASH1L related NDD ^36–43^. To avoid the contribution of potential differences in sleep behavior, sleep and wake states were analyzed separately. On a B6 background, *Ash1l^gt/+^* mice had a substantial increase in low-frequency delta oscillations (Figure 1J, and K, left panel). The increased delta power was restricted to the cortical electrode and not present in the hippocampal electrode. The increase in delta power in *Ash1l^gt/+^*mice occurred only during wakefulness. There was no change in any of the other frequency bands. CD-1 mutant mice had no changes in power spectral density during wake or sleep in the cortical or hippocampal electrodes (Figure 1K, right panel).

### ASH1L mutant mice have changes in sleep behavior

Sleep is a clinically relevant behavior for those with epilepsy and other NDD^44^. Thus, we studied how mutations in *Ash1l* affected sleep behaviors in mice. Because of interictal spiking during sleep in these mice, we wanted to control for possible sleep differences between WT and ASH1L mutant mice. Video EEG was scored for wake, non-REM sleep, and REM sleep. On both genetic backgrounds, there was no difference in the total amount of sleep or percentage of time spent asleep (Supplemental Figure 1A). On a B6 background, there was also no difference in the length or number of sleep bouts (Supplemental Figure 1B, left). However, on a CD-1 background, ASH1L mutant mice showed a gene dosage-dependent increase in the number of sleep bouts (Supplemental Figure 1B, right). In other words, their sleep was more fragmented, such that *Ash1l^gt/gt^* mice woke up almost twice as often throughout the day as their WT counterparts (Supplemental Figure 1C). The amount of REM sleep in ASH1L mutant mice was not different from WT on either mouse background. Furthermore, ASH1L mutant mice had intact circadian rhythms (Supplemental Figure 1D and E). It is possible that cortical seizures and spiking are triggering the frequent waking in mutant mice on the CD-1 background.

### Behavior in Ash1l^gt/+^ appears normal by initial screening

Based on the association of ASH1L with ASD and other behavioral diagnoses in humans, we tested whether ASH1L mutant mice for behavioral differences. To minimize variation, behavior was assayed only on the inbred B6 background, as per standard in the field. We did not perform behavioral studies on outbred mice on the CD-1 background. To see if *Ash1l^gt/+^*mice exhibited any changes in social behavior, we used a three-chamber social preference paradigm, in which the test mouse had the option of interacting with another mouse or an inanimate object (Supplemental Figure 1F, left). While WT mice prefer to interact with other mice, in some mouse models of ASD, this intrinsic social preference is impaired. However, *Ash1l^gt/+^* mice do not have a significant difference in social preference compared to WT mice (Supplemental Figure 1F, right). We also presented the test mice with a novel mouse or a familiar mouse (Supplemental Figure 1G). WT mice will generally have a preference for social novelty and spend more time interacting with the novel mouse. Both WT and mutant mice have intact preference for social novelty (Supplemental Figure 1G, right), indicating that short-term social memory is intact as well.

Although cognitive function can be difficult to measure in general, memory is one facet of cognitive ability that can be assessed in rodent models. To see if *Ash1l^gt/+^*mice had any memory deficits, we performed a novel object recognition test (Supplemental Figure 1H). When trained with two identical objects, WT mice prefer to interact with a novel object. *Ash1l^gt/+^* mice had intact preference for the novel object (Supplemental Figure 1H, right).

We tested whether ASH1L mutant mice were anxious or hyperactive, especially since either could alter results of other behavioral tests. As a prey species, mice naturally prefer to remain at the edges of any arena, and so time in the center of the arena is used as a measure of anxiety. *Ash1l^gt/+^* mice did not have a significant difference in the percentage of time spent in the center of the arena during the open field test (Supplemental Figure 1I, J left), suggesting that *Ash1l^gt/+^* mice are not overly anxious. *Ash1l^gt/+^*mice traveled the same distance as WT mice during the open field test, suggesting that *Ash1l^gt/+^* mice are not hyperactive (Supplemental Figure 1I, right). We assessed *Ash1l^gt/+^* mice for early gross motor deficits by performing a test of the righting reflex to see if pups could right themselves when placed in a supine position ^45^. Heterozygous *Ash1l^gt/+^* ^+^ and homozygous *Ash1l^gt/gt^* pups were able to right themselves normally (Supplemental Figure1K), suggesting that ASH1L mutant mice have grossly normal early motor development.

Because restricted and repetitive behaviors are a hallmark of ASD, we tested whether ASH1L mutant mice had any such behaviors. *Ash1l^gt/+^* mice did not exhibit increased self-grooming compared to WT counterparts (Supplemental Figure1L). Marble burying is another behavior that is thought to reflect repetitive or compulsive-like behaviors ^45^ (Supplemental Figure 1M). *Ash1l^gt/+^*mice did not have a significant difference in marble burying behavior (Supplemental Figure1M, top). Thus, this broad behavioral screen showed no major behavioral differences in *Ash1l^gt/+^* mice.

### ASH1L mutant mice have postnatal microcephaly

Motivated by the seizures in the mice, and the association of ASH1L with other NDD, we examined neuroanatomical phenotypes of ASH1L mutant mice that may underlie circuit changes. Although brain size and mass are indistinguishable at birth, *Ash1l^gt/gt^*mice on both genetic backgrounds have significantly decreased brain mass and cerebral area by the second postnatal week, persisting into adulthood (Figure 2A and B). This corresponds to a similar decrease in body mass, as well (Figure 2C). On a B6 background, adult *Ash1l^gt/+^* cerebrums had decreased length compared to WT brains, with no difference in cerebrum width (Figure 2D, middle panel). However, on a CD-1 background, *Ash1l^gt/+^* brains were increased in width relative to WT brains, while *Ash1l^gt/gt^* brains were decreased in length compared to WT (Figure 2D, right). Overall, loss of ASH1L leads to a decrease in postnatal brain growth that depends on both gene dosage and genetic background.

**Figure 2:**
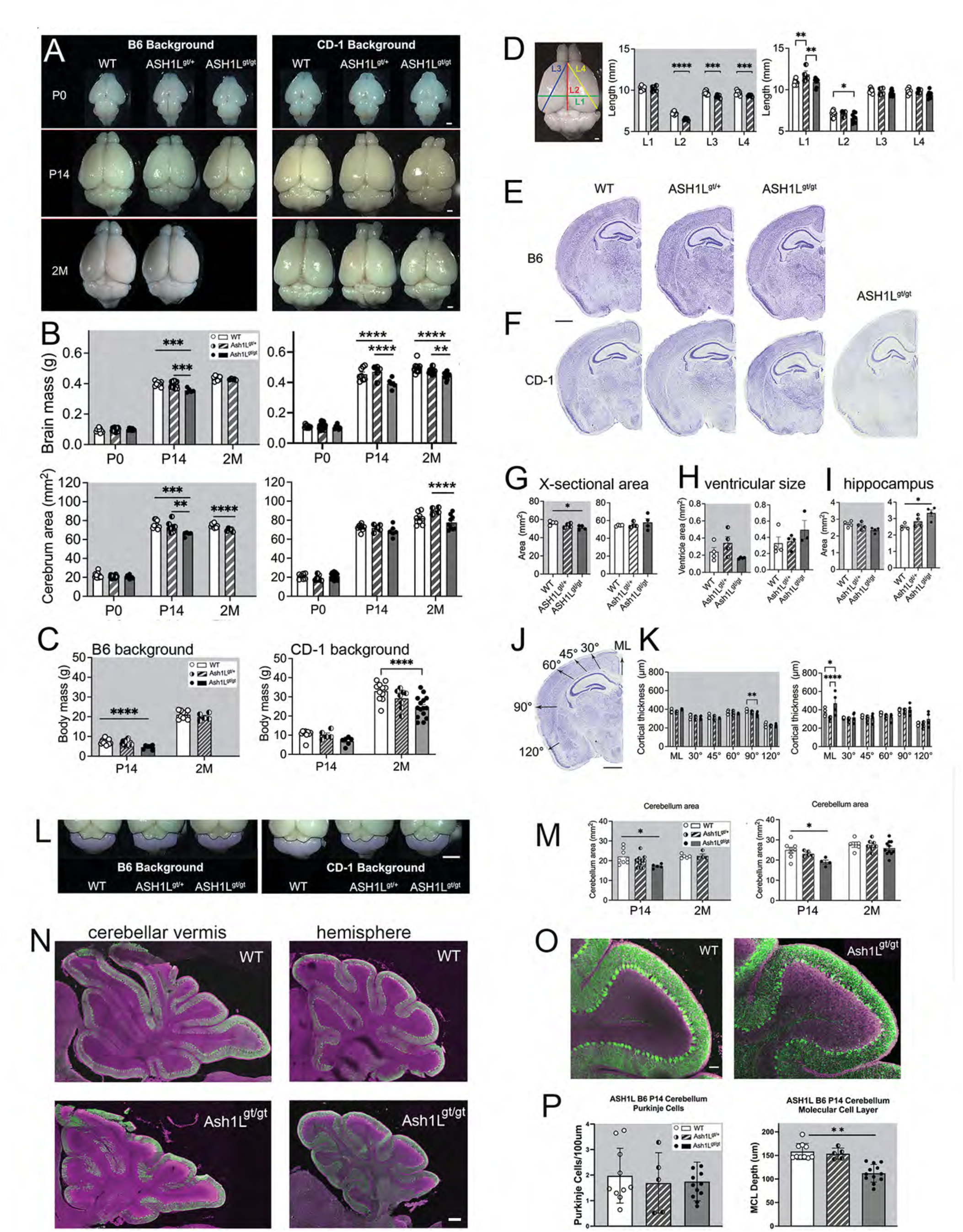
ASH1L mutant mice have postnatal microcephaly and cerebellar hypoplasia. (A) Representative images of mouse brains on a B6 (left) and CD-1 (right) genetic background at P0, P14, and 2M. Scale bar = 1mm. (B) On a B6 background, ASH1L mutant mice have a significant decrease in brain mass at P14. Data were analyzed with one-way ANOVA (F_2,25_ = 11.44, p = 0.0003) with Tukey’s post hoc multiple comparisons. At P0, a one-way ANOVA (F_2,29_ = 2.489, p = 0.1005) shows no significant difference in brain mass. At 2M, an unpaired t-test (t_12_ = 1.231, p = 0.2419) shows no significant difference between WT and *Ash1L^gt/+^* brain mass. N = 5-15 per genotype per time point. On a CD-1 background, ASH1L mutant mice have a significant decrease in brain mass at P14 and 2M. **** p <0.0001 by two-way ANOVA (age, genotype) with Tukey’s post hoc multiple comparisons shown. N = 4-28 per genotype per time point. On a B6 background, ASH1L mutant mice have a significant difference in cerebrum area. At P0, a one-way ANOVA (F_2,41_ = 1.679, p = 0.0662) shows no significant difference in brain area. At P14, a one-way ANOVA (F_2,22_= 9.367, p = 0.0011) with Tukey’s post hoc multiple comparisons shows a significant difference in brain area. At 2M, an unpaired t-test (t12 = 4.521, p = 0.0007) shows a significant decrease in *Ash1L^gt/+^* brain area. N = 5-19 per genotype per time point. On a CD-1 background, ASH1L mutant mice have a significant difference in cerebrum area. Two-way ANOVA (genotype, age) shows a significant effect of genotype (F_2,66_ = 8.401, p = 0.0006) and interaction between genotype and age (F_4,66_ = 2.684, p = 0.0389). Tukey’s post hoc multiple comparisons are shown. (C) On a B6 background, ASH1L mutant mice have a significant decrease in body mass at P14. Data were analyzed with one-way ANOVA (F_2,33_ = 13.29, p < 0.0001). At 2M, an unpaired t-test (t_13_ = 1.038, p = 0.3181) shows no significant difference between WT and *Ash1L^gt/+^* body mass. N = 5-16 per genotype per time point. Body mass is not significantly different at P0 (data not shown). On a CD-1 background, ASH1L mutant mice have a significant decrease in body mass. ** p = 0.0024 by two-way ANOVA (age, genotype) with Tukey’s post hoc multiple comparisons shown. N = 6-15 per genotype per time point. Body mass is not significantly different at P0 (data not shown). (D) Example of cerebrum lengths measured. Scale bar = 1 mm. On a B6 background, 2-month-old *Ash1L^gt/+^* mice have a significant decrease in brain length (L2, red line), the diagonal length of the left hemisphere (L3, blue line), and the diagonal length of the right hemisphere (L4, yellow line). N = 5-9 mice per genotype. Data analyzed by multiple unpaired t-tests with Welch correction. On a CD-1 background, 2-month-old ASH1L mutant mice have significant differences in brain width (L1, green line) and brain length (L2, red line). N = 4-9 mice per genotype. Error bars represent mean ± SEM. *P<.05, **P<.01, ***P<.001, ****P<.0001. (E, F) Representative images of Nissl-stained coronal brain sections at P14 on B6 (E) or CD-1 (F) background. CD-1 background shows one *Ash1L^gt/gt^* brain with microcephaly (left) and one *Ash1L^gt/gt^*brain with macrocephaly (right). Scale bar = 1mm. (G) On a B6 background, ASH1L mutant brains have a significant decrease in cross-sectional area by one-way ANOVA (F_2,9_ = 5.134, p = 0.0325). On a CD-1 background, ASH1L mutant brains have no significant difference in cross-sectional area (F_2,9_ = 0.5343, p = 0.6036) (H) There is no significant difference in ventricular size on the B6 background (F_2,9_ = 2.870, p = 0.1086) or on the CD-1 background (F_2,8_ = 1.221, p = 0.3446) by one-way ANOVA. (I) There is no difference in in terms of hippocampal area on a B6 background (F_2,9_ = 4.034, p = 0.0561). However, on the CD-1 background, ASH1L mutant mice have a significant increase in hippocampal area by one-way ANOVA (F_2,9_ = 7.091, p = 0.0142). (J) Schematic shows measurement of cortical thickness. Cortical thickness is measured at the midline (ML) and at various angles from dorsal to ventral. Method from Levy et al. 2021. Scale bar = 1mm. (K) On a B6 background, ASH1L mutant mice have significant differences in cortical thickness. Two-way ANOVA (angle, genotype) finds a significant effect of genotype (F_2,54_ = 3.523, p = 0.0365) and no significant interaction between angle and genotype (F_10,54_ = 1.438, p = 0.1893). Tukey’s post hoc multiple comparisons are shown. On a CD-1 background, ASH1L mutant mice have significant differences in cortical thickness. Two-way ANOVA (angle, genotype) finds a significant effect of genotype (F_2,54_ = 5.595, p = 0.0062) and interaction between angle and genotype (F_10,54_ = 2.082, p = 0.0422). Tukey’s post hoc multiple comparisons are shown. (L) Representative images of mouse brains on a B6 (left) and CD-1 (right) genetic background at P14 with cerebellar area that we measured overlayed in pink. Scale bar = 3 mm. (M) On a B6 background (left), cerebellum area is significantly decreased in ASH1L mutant mice at P14 by one-way ANOVA (F_2,21_ = 3.949, p = 0.0350), but not significantly different from WT at 2 months (t_7_ = 0.2235, p = 0.8295). On a CD-1 background (right), cerebellum area is significantly decreased in ASH1L mutant mice at P14 by one-way ANOVA (F_2,14_ = 5.087, p = 0.0218), but not significantly different from WT at 2 months (F_2,20_ = 0.7499, p = 0.4853). (N) In the B6 background at P14, staining for the Purkinje cell marker calbindin (green) and DAPI (magenta) at midline shows the cerebellar vermis and laterally shows the cerebellar hemispheres in both the WT and the mutant *Ash1L^gt/gt^* mouse. Scale bar is 500μm. (O) In the B6 background, the cerebellar lobes from WT and *Ash1L^gt/gt^* are immunostained with the Purkinje cell marker calbindin (green) and DAPI (magenta). Scale bar is 100μm. (P) Quantification of Purkinje cell numbers shows no statistically significant decrease ithe B6 background. However, measurement of the molecular layer shows a decrease in thickness in the *Ash1L^gt/gt^* mouse (F_2,24_=22.67, p<0.0001) by one-way ANOVA.

Because ASH1L mutant mice have postnatal microcephaly, we sought to determine whether specific brain regions are affected. We examined mouse brain sections at P14, when microcephaly is present. On a B6 background, Nissl-stained coronal sections from ASH1L mutant mice had decreased cross-sectional area (Figure 2E, top and 2F, left panel). However, on a CD-1 background, while some brains had decreased cross-sectional area, others had increased cross-sectional area (Figure 2E, bottom and 2F, right). There were no significant differences in size of the lateral ventricles, ie no hydrocephalus(Figure 2G). However, ASH1L mutant mice on a CD-1 background have a significant increase in hippocampal size (Figure 2H). To see if the changes in brain size could be attributed to differences in cortical development, we measured cortical thickness (distance from the pial surface to the corpus callosum) at varying angles (Levy et al. 2021^46^ (Figure 2J). ASH1L mutant brains had subtle changes in cortical thickness, laterally in B6 mutant mice (Figure 2K, left) and at the midline in CD-1 mutant mice (Figure 2K, right). To summarize, no single brain area emerges as the driver of microcephaly in ASH1L mutant mice. Instead, most likely, small changes in various brain areas add up to an overall decrease in brain volume.

Microcephaly in ASH1L mutant mice could arise from a decrease in cell number. Numerous lines of evidence suggest that ASH1L could affect cell proliferation. Generally, Polycomb / Trithorax signaling affects cell cycle progression.^47–49^ In the mouse brain, ASH1L is expressed at the right time and place to affect neurogenesis, as it is enriched in the embryonic ventricular zone and subventricular zone.^50^ Although the effect of ASH1L deletion on progenitor cell fate in the brain *in* vivo has been demonstrated^51^, no difference in progenitor proliferation was observed in our mouse lines with hypomorphic mutations.

In our hands, analysis of distribution and density of cells at P14 is also unchanged on either B6 or CD-1 background, so that the mild microcephaly might not be from impaired neurogenesis or gliogenesis (Supplemental Figure 2A, B). However, we examined the cortex at postnatal day 0 to determine whether ASH1L mutant mice had any differences in cell number, reflecting antenatal proliferation defects (Supplemental Figure 2C-G). Cell fate changes or premature depletion of neural progenitor cells in ASH1L mutant mice could manifest as a decrease in the total number of later born neurons which are located in the upper-layers of the cortex ^52^. However, at birth, *Ash1l^gt/gt^* mice lack differences in the total number of all cells (Supplemental Figure 2F), neurons expressing the upper layer marker SATB2 (Supplemental Figure 2C), and neurons expressing the lower layer markers Ctip2 (Supplemental Figure 2D),) and Tbr1 (Supplemental Figure 2E).

Many NDD-associated genes cause defects in neuronal migration.^53–57^ Even if there is no difference in the total number of upper- and lower-layer neurons, ASH1L mutations could prevent neurons from migrating to the correct location. To determine whether ASH1L mutations affect neuronal migration, we compared the vertical distribution (from the pial surface to the ventricular zone) of upper- and lower-layer neurons between *Ash1l^gt/gt^* and WT mice. At birth, there is no change in the vertical distribution of neurons positive for the upper layer marker SATB2 or the lower layer markers Ctip2 and Tbr1, suggesting that loss of ASH1L does not cause a neuronal migration defect. The distribution of all DAPI+ nuclei is also unchanged. Overall, *Ash1l^gt/gt^* mice do not appear to have altered embryonic development of the cortex, either neuronal proliferation or neuronal migration.

### ASH1L mutations cause changes in the cerebellum

While we focused on the cortex and hippocampus initially because of our interest in the neurocognitive phenotypes, other brain regions are affected by ASH1L. In both backgrounds, at P14, the gross area of the cerebellum is statistically decreased in the *Ash1l^gt/g^*^t^ genotypes (Figure. 2L and M). Sagittal sections at the midline showing the vermis and and laterally, showing the cerebellar hemispheres demonstrate hypoplasia in the *Ash1l^gt/g^*^t^ cerebellum in both regions in B6 mice (Figure 2N). Analysis of higher power images of the lobes demonstrated no change numbers of Purkinje neurons. However, disruption of ASH1L results in a thinner molecular layer. The thinner molecular layer may be indicative impairment of dendritogenesis of Purkinje neurons, consistent with the dendritic phenotype previously observed in human experimental systems.

### ASH1L mutant mice have decreased arborization of layer V pyramidal cells

Because ASH1L mutant mice showed no consistent differences in brain region areas or in neuronal numbers, postnatal microcephaly in ASH1L mutant mice could arise from a failure of neurite outgrowth. We tested whether ASH1L would also affect neurite outgrowth *in vivo*. In the course of performing whole cell patch clamp electrophysiology, we were able to visualize biocytin filled neurons age p24-27 (Figure 3A). We confirmed their position in CA1 of the hippocampus (Figure 3B) and overlaid images with concentric circles for Sholl analysis (Figure 3C). The number of crossings showed defects in both apical and basal dendrites. In order to perform the study in both backgrounds, BL6 and CD-1, Golgi staining was performed to visualize neurons in adult mice. Because previous work had examined deep layer cortical excitatory neurons^27^, we analyzed layer V pyramidal cells (Figure 3E-F). Across both genetic backgrounds, ASH1L mutant neurons had decreased arborization of basal dendrites by Sholl analysis (Figure 3G, H). This decrease was greatest in the middle of the neurites, less so proximally and distally. On a B6 background, *Ash1l^gt/+^* neurons did not have a significant difference in number of branch points, neurite length, basal complexity index, or soma area (Figure 3I-L, left panels). On a CD-1 background, while soma area and number of branch points were still unchanged, *Ash1l^gt/+^* mutant neurons had decreased basal complexity index and decreased neurite length (Figure 3I-L, right panels). Thus, microcephaly in the ASH1L mutant mouse could be, at least in part, due to a decreased neurite outgrowth. This finding fits with previous work showing that loss of ASH1L may decrease neuronal responsiveness to growth factor signaling.^27^

**Figure 3:**
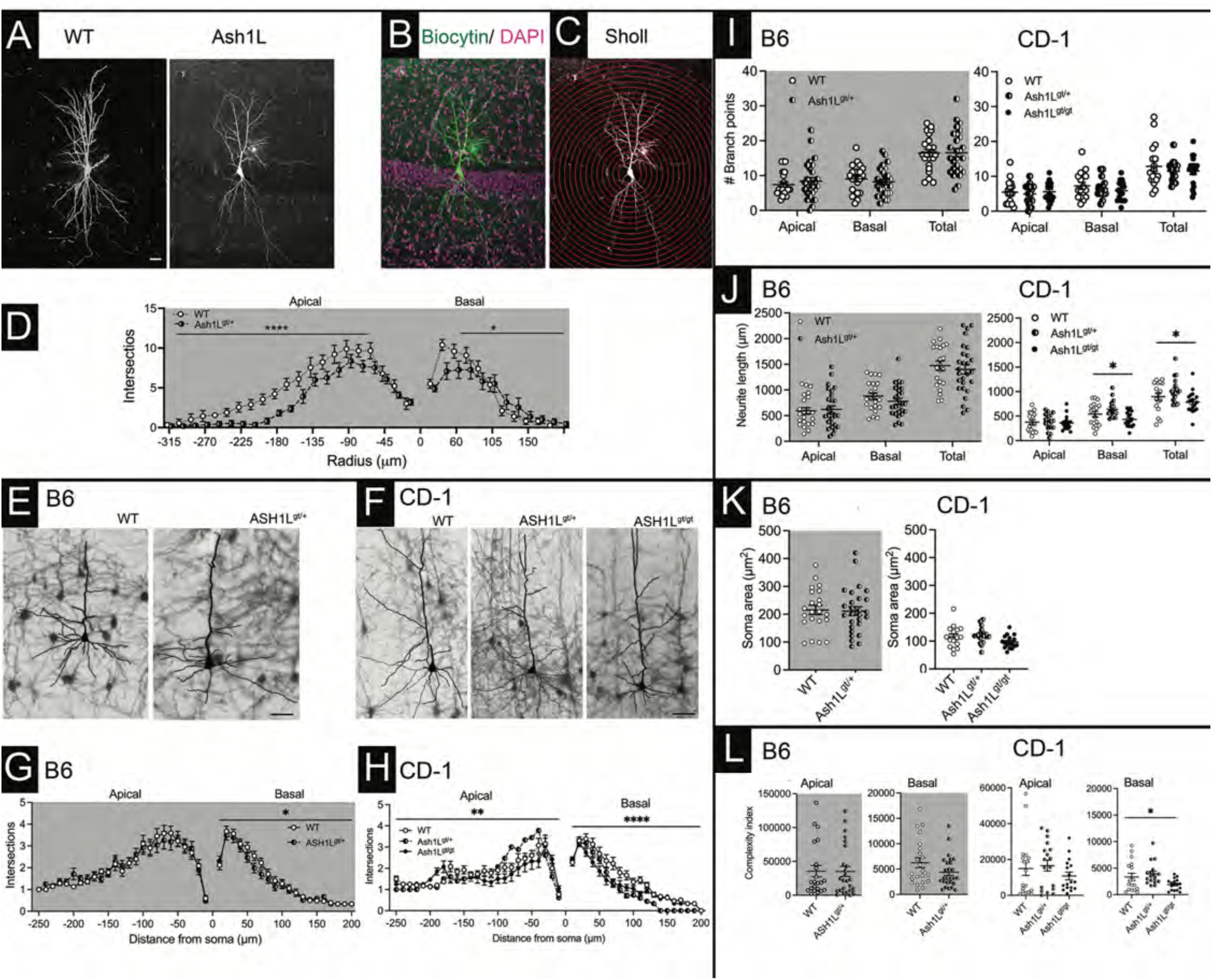
Dendritic defects in Ash1L mutants. A) Biocytin-filled neurons in the CA1 region of the hippocampus in WT and *Ash1L^gt/+^* mice at postnatal day 23-25. (B) DAPI counterstain to verify location of *Ash1L^gt/+^* neuron in CA1 (C) Concentric circle overlay for Sholl analysis with radius increasing in 15μm intervals. Scale bare is 30 μm. (D) On a B6 background, *Ash1L^gt/+^* neurons have decreased intersections of the basal dendrites by Sholl analysis. ASH1L mutant neurons have a significant difference in the number of intersections of apical and basal dendrites. Apical: **** p = 0.0001 by two-way ANOVA (genotype, distance from soma). Basal: * p < 0.039 by two-way ANOVA (genotype, distance from soma). Representative images of layer V pyramidal cells on B6 (E) and CD-1 (F) backgrounds visualized with Golgi stain. Scale bar = 50 µm. (G) On a B6 background, *Ash1L^gt/+^* neurons have decreased intersections of the basal dendrites by Sholl analysis. *p = 0.025 by two-way ANOVA (genotype, distance from soma). (H) On a CD-1 background, ASH1L mutant neurons have a significant difference in the number of intersections of apical and basal dendrites by Sholl analysis. Apical: ** p = 0.0022 by two-way ANOVA (genotype, distance from soma). Basal: **** p < 0.0001 by two-way ANOVA (genotype, distance from soma). (I) On a B6 background, ASH1L mutant neurons have no significant difference in the number of apical (t_47_ = 0.7221, p = 0.4738), basal (t_47_ = 0.7523, p = 0.4556), and total (t_47_ = 0.0607, p = 0.9519) branch points by unpaired t-test. On a CD-1 background, ASH1L mutant neurons have no significant difference in the number of apical (F_2,48_ = 0.2147, p = 0.8076), basal (F_2,49_ = 0.7994, p = 0.4554), and total (F_2,51_ = 0.5308, p = 0.5914) branch points by one-way ANOVA. (J) On a B6 background, ASH1L mutant neurons have no significant difference in apical (t_47_ = 0.3156, p = 0.7537), basal (t_47_ = 1.194, p = 0.2385), and total (t_47_ = 0.5521, p = 0.5835) neurite length by unpaired t-test. On a CD-1 background, ASH1L mutant neurons have no significant difference in apical neurite length. However, basal and total neurite length differ by genotype. * p < 0.05 by Kruskal-Wallis test. (K) On a B6 background, ASH1L mutant neurons have no significant difference in soma area by unpaired t-test (t_47_ = 0.1756, p = 0.8614). On a CD-1 background, ASH1L mutant neurons have no significant difference in soma area by one-way ANOVA (F_2,48_ = 1.609, p = 0.0635). (L) On a B6 background, ASH1L mutant neurons have no significant difference in apical or basal complexity index by Mann-Whitney test. On a CD-1 background, basal neurites show a significant difference in complexity index, * p = 0.0107 by Kruskal-Wallis test. B6: N = 22-27 cells per genotype from 6 mice per genotype (3 males, 3 females); CD-1: N = 18 cells per genotype from 6 mice per genotype (3 males, 3 females).

### ASH1L mutant mice have changes in inhibitory interneurons

ASH1L is expressed in both excitatory neurons and inhibitory neurons,^23–26^ so mutations in ASH1L could affect inhibitory neurons in cell-autonomous and non-cell-autonomous ways. Also, inhibitory interneuron dysfunction is common in epilepsy. Therefore, examined populations of inhibitory interneurons in ASH1L mutant mice. We quantified inhibitory interneurons expressing somatostatin (SST). On a B6 background, ASH1L mutant mice have a significant decrease in SST+ cells in the hippocampus, particularly in hippocampal CA1 (Supplemental Figure 3A, and B, top). Conversely, on a CD-1 background, ASH1L mutant mice have an increase in SST+ cells in the hippocampus and hippocampal CA1 (Supplemental Figure 3B, bottom panel). We also quantified the most abundant of the inhibitory neurons, parvalbumin (PV)-expressing cells.^58^ On both genetic backgrounds, ASH1L mutant mice had no difference in the number of PV+ cells, either in the cortex or hippocampus (Supplemental Figure 3C and D).

Although there is no difference in the number of PV+ cells, we tested if more subtle changes in inhibitory signaling could be detected. PV+ basket cells make perisomatic synapses onto the somas of neocortical pyramidal cells.^58^ We quantified the number of perisomatic puncta from PV+ terminals onto NeuN+ layer V pyramidal cells as a correlate of perisomatic synapses. Across both genetic backgrounds, there is no significant difference in the number of perisomatic puncta between WT and *Ash1l^gt/gt^*mice (Supplemental Figure 3E-J). Surprisingly, on both genetic backgrounds, the heterozygous *Ash1l^gt/+^* mice differed significantly from WT and *Ash1l^gt/gt^* mice, suggesting an overdominant phenotype (Supplemental Figure 3F, I). However, the polarity of the difference was background dependent. On a B6 background, heterozygotes had more perisomatic puncta than WT and *Ash1l^gt/gt^* counterparts (Supplemental Figure 3E-G). Conversely, on a CD-1 background, heterozygotes had more perisomatic puncta than WT and *Ash1l^gt/gt^*counterparts (Supplemental Figure 3H-J). For example, on a CD-1 background, 92% of *Ash1l^gt/+^* layer V pyramidal neurons had 10 or fewer perisomatic puncta, compared to 62-67% of WT and ASH1L^gt/gt^ neurons.

It is unclear what mechanisms drive this overdominant and background-dependent phenotype. The gene dosage-dependent difference could represent an adaptation in *Ash1l^gt/+^* mice that *Ash1l^gt/gt^* mice are unable to execute. Alternatively, if ASH1L mutations are acting in a dominant negative manner, the heterozygous mutant mouse would be expected to have a more severe phenotype than the homozygous mutant. Nevertheless, the same pattern does not appear for other phenotypes assayed. Nor are we aware of this pattern in other mouse models. It is also not clear whether other types of inhibition are increased or decreased to make up for a relative decrease or increase in perisomatic inhibition by PV+ basket cells. Notably, WT mice on a CD-1 background appeared to have more perisomatic synapses than B6 WT mice (median of 5 puncta/soma in WT B6 mice versus a median of 9 puncta/soma in WT CD-1 mice). WT CD-1 mice also appear to have more PV+ neurons in the cortex than WT B6 mice (Supplemental Figure 3F, I). However, PV+ interneurons have not been specifically studied on a CD-1 genetic background.

### Patients with ASH1L have a diversity of genetic mutations

In order to understand the human genetic and phenotypic diversity, we conducted a clinical phenotyping study. For our evaluation, we conducted medical history and records survey, medical/neurological examination, a neuropsychological evaluation, and a standard EEG.

We found that patients had a diversity of mutations in our study including missense and frameshift mutations throughout the length of the ASH1L coding sequence. Overall, we were able to evaluate 10 male patients and 13 female patients ranging from 3 years of age to 32 years of age at the time of the initial evaluation (Table S1, the supplement to Figure 4). Mutations in ASH1L were both inherited and *de novo* (Table S1). However, some were unable to be determined because biological parents were not available sequenced. Both male and female patients had splice site (a male and female sibling pair), nonsense, missense, and frameshift mutations (Figure 4A). While we had 23 total patients with 23 distinct mutations since one participant had 2 missense mutations, and a sibling pair had the same mutation (Table S1, Figure 4A, B, C). The majority of the cohort (16/23) had mutations that could cause early termination of translation (nonsense and frameshift) (Figure 4A, D), but may actually result in loss of function through nonsense mediated decay of mRNA. Despite the relatively small number of patients in the initial cohort, we were able to observe a variety of phenotypes.

**Figure 4:**
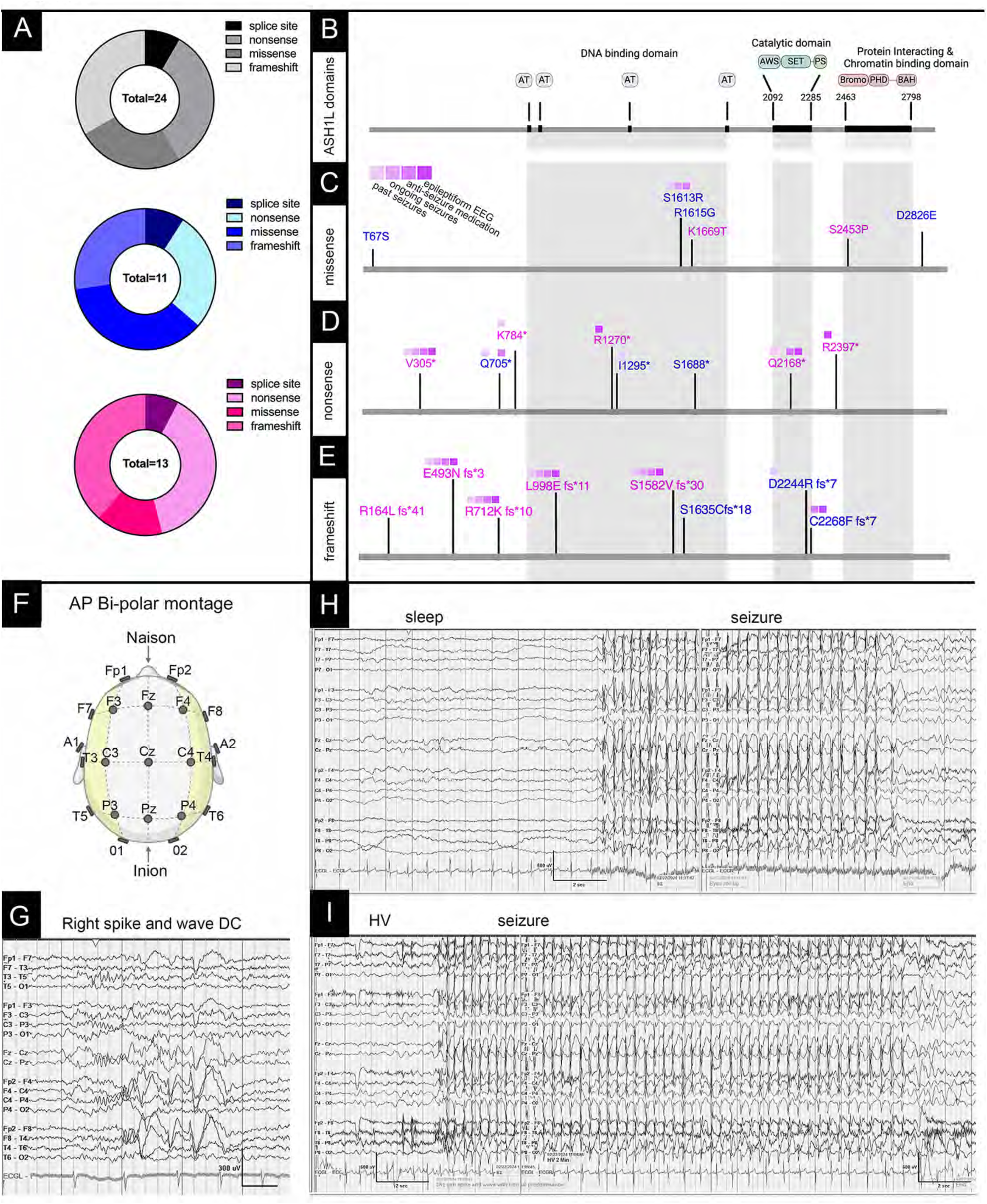
Genetics and epilepsy in ASH1L: (A) We performed in-person evaluations of 23 total patients with mutations in ASH1L. The type of mutations in ASH1L were abstracted from the clinical genetic reports: 2 splice site, 8 frameshift, and 8 nonsense, and 6 missense mutations. A total of 24 mutations are present since one subject has two separate missense mutations on the same copy of the gene (top). Of the 10 boys (blue, middle), there are 11 total mutations with 1 subject with a splice site mutation, 3 each with frameshift and nonsense mutations and 4 missense mutations within 3 subjects. In 13 girls (pink, bottom), the mutations were distributed in roughly the same proportions with 1 mutation affecting splicing, 5 each for frameshift and nonsense mutations, and finally 2 missense mutations. (B) The ASH1L protein with functional domains that are shown for reference for position of (C) missense mutations and (D) nonsense mutations and (E) frameshift mutations. For panels C-E mutations that are represented in blue are in boys and those represented in pink are in boys. Epilepsy burden for each patient is represented in purple including past and ongoing epilepsy, anti-seizure medications, and epileptiform EEG from our in-person study. (F) Electroencephalograms (EEGs) were performed on subjects in the study and studies are shown in the anterior/ posterior bi-polar montage. (G) Focal epileptiform discharges (right sided 3Hz spike and slow waves) are observed, as well as generalized epilepsy where (H) shows a generalized seizure from sleep and (I) shows another generalized seizure provoked by hyperventilation (HV).

### ASH1L mutations can cause epilepsy and other neurological symptoms

The constellation of brain phenotypes reported for ASH1L patients may result in a variety of brain phenotypes. Affected individuals had neurological and neuropsychiatric disorders including epilepsy, deveopmental delay, autism spectrum disorder, intellectual disability, attention deficit hyperactivity disorder, obsessive compulsive disorder, and tic disorder. Based on analysis of medical history and the results of the testing, it appeared that there was a sexually dimorphic presence of epilepsy and autism spectrum disorder (Table S1), while other neuropsychiatric disorders were present in both sexes at similar rates.

Case reports suggest that about one-third to two-thirds of children with ASH1L mutations have seizures^9^. However, more in-depth characterization has not been available. In the girls, 7 of the 13 participants had a history of clinical epilepsy, mostly generalized non-motor (absence), with occasional generalized tonic-clonic seizures. Of the girls, nine of the thirteen had epileptiform activity on the EEGs performed in our study. Many of the EEG’s showed occasional bursts of generalized polyspike/spike and slow wave discharges. Asymmetry was common as well, with right sided spike and slow waves (Figure 4E, F). One patient had seizures on a baseline EEG performed for our study (Figure 4G). Furthermore, hyperventilation in that patient caused an 18-second seizure with behavioral correlates of eye fluttering and eyes rolling upward (Figure 4H). Two female participants had a history of photo-stimulated epilepsy, and photic driving was not performed in our EEG laboratory on studies with those patients. In contrast, of the male patients, one had encephalopathy on EEG, and every other participant had a normal baseline EEG. Only one boy out of ten demonstrated epileptiform activity by EEG and only during hyperventilation. Indeed, three of the nine (1/3) had a history of seizures with only the youngest of the male participants on anti-convulsant medication, with currently active epilepsy. Other neurological issues include history of hypotonia, eye movement abnormalities, and mild ataxia on neurological exam that was consistent with the cerebellar phenotype observed in mice (Figure 2L-P). Neuroimaging was often normal, but included hypomyelination and also mild Chiari malformations (Table S1).

### ASH1L patients have developmental delay and other medical symptoms

In our patient cohort we observed no statistical difference with regard to biometrics between male and female participants. There were no statistically significant differences in height, weight, or head circumference in our cohort (Figure 5A). However, most children with ASH1L mutations were late meeting at least one developmental milestone (Figure 5B). While most children have developmental delay, one boy experienced regression, which was also observed in one published case ^2^. Overall, both motor and speech development are delayed in children with ASH1L mutations. Boys appeared to be more severely affected than girls in both motor and language domains, as there was a statistically significant difference in the age at first steps (Figure 5B, right).

**Figure 5:**
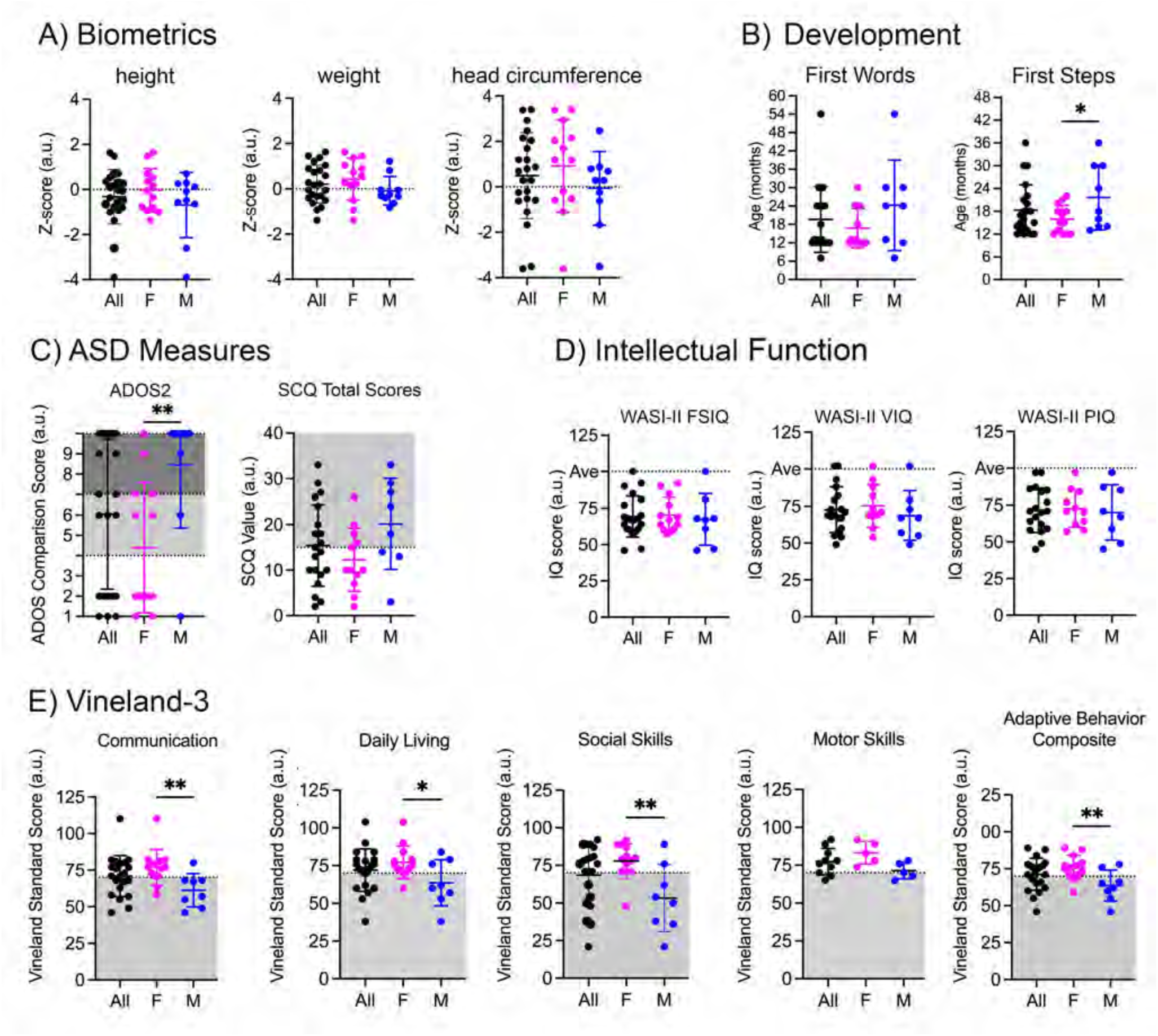
Sex differences in in patients with ASH1L syndrome: We compared subjects with ASH1L mutations with respect to gender. The entire cohort is shown in black. Measures taken in girls are pink and blue represents values in boys. The mean of each data set is shown with error bars which are standard deviation. (A) We measured height, weight and head circumference in all 23 subjects and there were no statistical differences between the boys and girls, although the mean of height is below average in boys and head circumference is above average in girls. (unpaired t-test with Welch’s correction, p= 0.22 for height, p= 0.14, p= 0.087 for head circumference) (B) We analyzed developmental milestones in boys and girls. Boys were significantly more delayed than girls for age of first steps. In terms of development, many kids are outside of normal milestone acquisition time frame (10-14 months, light green). For age of first words, only 8 of the 10 boys were included because two of the boys were non-verbal at the time of the evaluation. 9 of 10 boys were included in the analysis because the youngest subject was not yet walking at the time of the evaluation. (unpaired t-test, p=0.123 for first words and p=0.0427 for first steps). (C) Autism spectrum disorder is evaluated by the Autism Diagnostic Observation Schedule-2 ADOS2, an instrument using direct observation by a trained observer and by the Social Communication Questionnaire (SCQ), a parental rating scale. For ADOS2 (left) subjects had scores in mildly range (white region on the graph), in the moderate range (light gray), and severe (the dark gray). All 13 of the girls in our study and 9 boys were assessed and the boys were statistically more severely affected than the girls (unpaired t-test with Welch’s correction, p=0.008). In terms of the SCQ (right), scores over 15 are considered abnormal and the average of the boys scores fall within the abnormal range. We have scores for 12 girls and 8 boys. The p-value of the unpaired t-Test is for comparing scores between boys and girls is 0.051. (D) In order to assess for intellectual disability, our members of our cohort within the appropriate age ranges (13 girls and 8 boys) underwent testing by the Weschler Abbreviated Scale of Intelligence, Second Edition (WASI-II). Full Scale Intelligence Quotient (FSIQ), Verbal Intelligence Quotient (VIQ), and Performance Intelligence Quotient (PIQ) are shown. For FSIQ the average of the cohort (69.09) is just within the range of mild intellectual disability (50-70) without significant differences between male and female subjects (unpaired t-Test, p=0.23). Results of VIQ and PIQ are similar suggesting no specific domain impairment. Means of VIQ and PIQ are 72.5 and 71.5, respectively. There are also no significant differences between male and female subjects for VIQ (with p=0.36) and PIQ (with p=0.716). (E) The Vineland is designed to measure adaptive behavior (ability to live independently) of individuals. We show 4 domains: Communication, Daily Living Skills, Socialization and Motor Skills. The domain scores yield an adaptive behavior composite (far right panel). The Vineland is given to all 23 participants, but only participants under the age of 9 years old take part in the motor skills assessments. Score falling below 70 (in the gray zone) indicate significant skills deficit below similarly aged peers. Males were significantly more affected than females in the cohort in domains, which then translated to a significantly lower composite score, as well (unpaired t-test, p= 0.09 for Communication, p= 0.027 for Daily Living Skills, p= 0.003 for Social Skills, p= 0.02 for Motor Skills, and p=0.007 for the Adaptive Behavior Composite).

However, since ASH1L is ubiquitously expressed, organ systems other than the brain may be affected (Table S1). Overall, while congenital cardiac anomalies are more common in children with ASH1L mutations than in the general population, but they are still relatively rare and of unknown clinical significance with only one participant in our group who was affected. Other organ systems are also affected. In our cohort, there were two instances of precocious puberty, that was not consistent with other family members (first degree relatives). However, the most prevalent non-brain phenotypes are in the gastrointestinal system with most participants having upper, lower, or hepatic abnormalities. Tongue tie, and swallowing are common struggles, with participants endorsing dysphagia. Multiple participants in our study, as well as reports in the literature ^11^, report constipation (Table S1). The increased prevalence of GI disorders could reflect a multitude of changes, possibly developmental, or in the immune system, microbiome, intestinal permeability, motility, secretion, behaviors, or alterations in the enteric nervous system.

### ASH1L mutations can cause autism spectrum disorders predominantly in males and other neuropsychiatric disorders across sexes

In contrast to the female cohort of ASH1L affected children, autism spectrum disorder phenotype was more penetrant in the male cohort. Of the 9 male participants, n=6 had a clinical ASD diagnosis confirmed with ADOS and n = 2 had either clinical ASD diagnosis or positive ADOS. Only the youngest participant (3 years old) did not have a diagnosis of ASD or positive ADOS (Table S1). For the others, there was congruence of ADOS with parental rating scales. Additionally, one of the male affected subjects experienced a prior history of developmental regression. In contrast, of the female cohort, only 3 of the 13 had a clinical ASD diagnosis that was confirmed with the ADOS. Another parental rating scale, the Social Responsiveness Scale (SRS-2) also showed greater impairment in boys vs. girls in terms of social awareness, but not in other SRS-2 subdomains (Supplement to Figure 5A, B). In contrast to autism and epilepsy, ASH1L mutations cause intellectual disability to roughly the same degree across both sexes without dimorphism as measured via overall, verbal, and visuospatial intelligence (Figure 5D). In addition, irritability (Supplement to Figure 5C), ADHD or hyperactivity, (Table S1 and Supplement to Figure 5D, anxiety (Table S1) and obsessive-compulsive phenotypes (Table S1) are also present across the board without sexual dimorphism. Overall, these symptoms are reflected in the Vineland-3, a measure of adaptive behavior or the collection of skills needed to function independently in daily life, with boys being more severely affected than the girls, especially in domains affecting social skills and communication (Figure 5E).

### Sexually dimorphic circuit effects of ASH1L inactivation

Sex differences in expression of autism and epilepsy/ EEG abnormalities suggest differential effects of sex on specific circuits caused by ASH1L deficiency. While we observed differences in the rate of autism and epilepsy in males and females, whether these were a result of a sex difference in the effect of ASH1L insufficiency is unknown since we are studying a relatively small number of affected individuals. Our cohort contains a larger number of female probands with early truncations in contrast to males, so that the nature of the mutation might underlie the apparent differences. To better understand this difference, we returned to our mouse model where we could determine whether a sex-specific effect might still be present in specific cells or circuits. We performed whole-cell patch clamp electrophysiology on CA1 pyramidal neurons in the hippocampus of male and female mice between postnatal day 23-26 (Figure 6), choosing this brain region because seizures arise frequently from the hippocampus^59^ (Figure 1).

**Figure 6:**
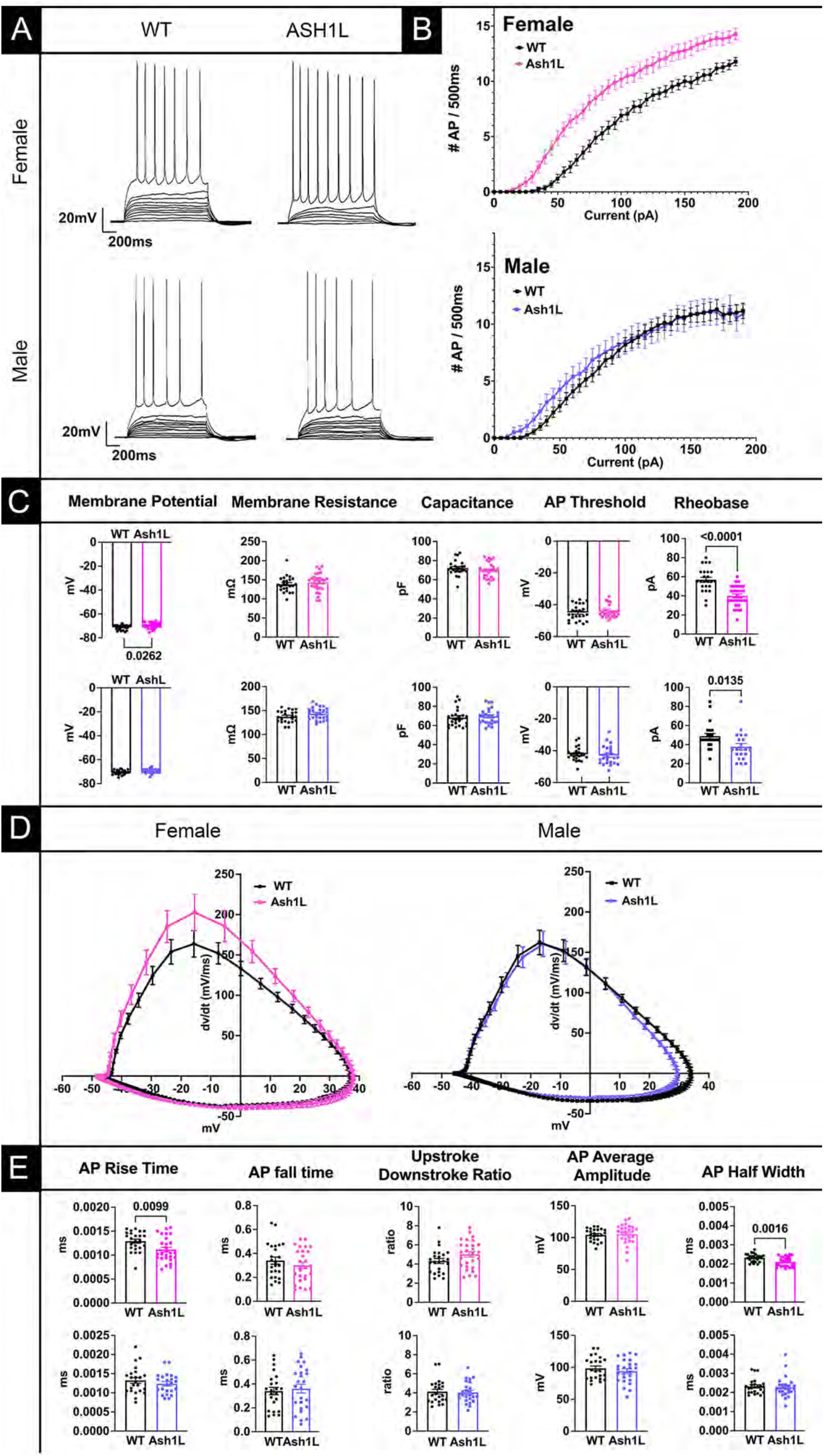
Hyperexcitability in Action Potential Firing from Female Hippocampal CA1 Pyramidal Neurons: (A) Action Potentials (AP) are triggered by stepwise current injections ranging from -20 to 195 and increasing in 5pA steps. (Top) Neurons from *Ash1L^gt/+^* female mice fire action potentials with lower amounts of current than WT female neurons. In contrast, (bottom) male wild type and *Ash1L^gt/+^*neuron show few differences. (B) Input/Output curves show clear separation between female WT and *Ash1L^gt/+^* neuronal AP firing in response to a range of current injection (top, magenta) in contrast to male neurons (bottom, blue). (C) Intrinsic properties are show for female (top) and male (bottom) wild type and *Ash1L^gt/+^* mice: membrane potential, membrane resistance, capacitance, AP threshold potential, and Rheobase (minimum current injection for first AP). (D) Phase plot analysis of AP in wildtype and *Ash1L^gt/+^* female (left) and male (right) shows differences in the rate of membrane potential depolarization in female neurons in contrast to male neurons. (E) Analysis of AP properties in neurons from female (top) and male (bottom): AP rise time, fall time, ratio of upstroke to downstroke, amplitude, and half width. Mice were between post-natal day 23-26 for both male and female mice. n= 7 WT female mice and 24 cells. n= 7 for *Ash1L^gt/+^* female mice, and 27 cells. n= 7 for WT male mice and 23 cells. n= 7 for male *Ash1L^gt/+^* mice and 26 cells. Data are represented as mean+/ SEM. p values were determined by unpaired t-tests and significant p values less than 0.05 are shown on graphs.

ASH1L deficiency modifies the excitability and the action potential shape of the hippocampal CA1 principal neurons in female mice alone. We performed our experiments in current clamp mode (Figure 6). We injected current in a step-wise manner (Figure 6A and 6B) and measured intrinsic properties. Intrinsic properties (membrane resistance, capacitance, and threshold voltages) were unchanged in neurons from both male and female mice (Figure 6C). However, resting membrane potential was slightly, but significantly, elevated in female mice. In terms of rheobase (lowest current causing action potential firing), neurons from both male and female mice showed significant decreases over control neurons. However, the extent of the difference is greater in neurons from female mice (Figure 6C).

Analysis of action potentials in female mice demonstrates significant differences in female vs. male mice. The changes in the action potentials are reflected in phase plots (first derivative of membrane voltage/ over time) (Figure 6D), where the depolarization phase in *Ash1L^gt/+^* female neurons is significantly different from that of controls, whereas no changes are observed in male *Ash1L^gt/+^* male neurons. Analysis of action potential characteristics (Figure 6E) show a decrease in the AP half-width, likely due to the decreased rise time, which is also significantly decreased. The peak amplitude and upstroke/ downstroke ratio are unchanged. In contrast, none of these parameters are significantly altered in male mice with *Ash1l ^gt/+^* compared to male control animals. Thus, the response to current injection overall demonstrates a hyperexcitability present in neurons from female mice with a shift of the input/ output curve that is not apparent in neurons from male mice.

To determine whether the same changes are observed in other brain circuits, we also performed the same studies in the cortex (Supplement to Figure 6). In contrast to the hippocampus, no changes are observed in layer 5 pyramidal neurons in the somatosensory cortex in female mice. Thus, the sex effect for Ash1L may be circuit specific. Furthermore, the changes in some specific circuits but not others may underlie the sexually dimorphic presentation of ASH1L deficiency in humans.

### ASH1L regulates transcriptional programs important for neural circuit development in a sex-specific manner in mouse hippocampi

We posit that the sex-specific electrophysiological phenotypes observed in the hippocampus of Ash1l mutant mice could be underlie by distinct molecular programs in males and females driven by Ash1l dysfunction. We performed RNA sequencing (RNA-seq) in wild type (WT) and heterozygous (HET) *Ash1L^gt/+^* hippocampi isolated from P26 male and female mice to match with electrophysiological experiments. To assess sex-specific effects, we first conducted separate comparisons for HET (*Ash1L^gt/+^*) vs. WT males and for *Ash1L^gt/+^* vs. WT females. Analysis of *Ash1L^gt/+^* vs. WT comparisons identified 176 differentially expressed genes (DEGs) in females and 157 DEGs in males (fold change > 1.2, FDR < 0.05). Only 1% of DEGs overlapped between the two datasets, with just seven shared genes (Cd40, Fbxl9, Msantd7, Or1f19, Rpl18a, Trpm7 (Fig. 7A-C, Supplement to Figure 7, and Table S2). Gene set enrichment analysis (GSEA) which considers fold change and statistical significance of the DEGs revealed distinct biological pathways affected in each sex. By analyzing cellular component in female *Ash1L^gt/+^* mice, genes involved in myelin sheath formation, ribosomal subunits, mitochondria, and the proteasomal complex were downregulated compared to WT females. In contrast, male HET *Ash1L^gt/+^* mice showed downregulation of neuronal spine and axon-associated genes, while mitochondrial protein encoding genes were upregulated compared to WT males. Additionally, analysis of biological process, showed male and female specific signatures (Figure 7E-G, Supplement to Figure 7, Table S3). *Ash1L^gt/+^* female compared to WT females showed enrichment for genes involved in sensory perception among the upregulated genes and downregulation of genes involved in metabolic pathways. In contrast, *Ash1L^gt/+^*males compared to WT males had an overrepresentation of genes involved in neural circuit development and function. We observed upregulation of genes important for glial differentiation, telencephalon development, as well as axon guidance, and downregulation of genes involved in synaptic plasticity, and neurogenesis (Supplement to Figure 7, Table S3).

**Figure 7:**
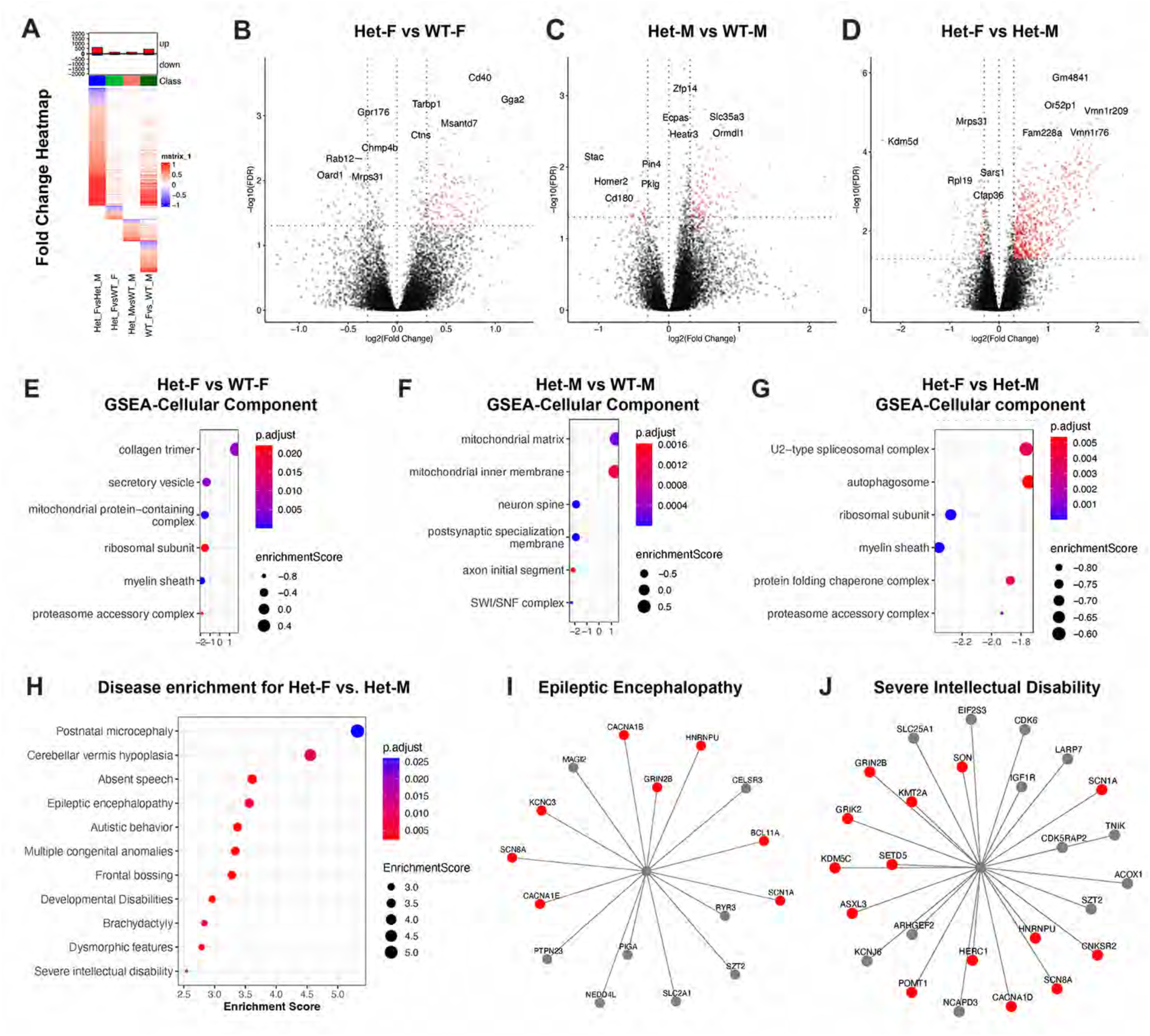
ASH1L regulates gene programs essential for neural circuit development in a sex-specific manner. (A) Heatmap shows DEGs for all comparisons between Heterozygous (HET) and wild type (WT) females and males. The total number of DEGs are listed. (B-D) Volcano plots showing DEGs above adjusted p value of 0.05 (red dots) that are either downregulated (left side of central dotted lane) or upregulated (right side of central dotted lane). The top 10 DEGs are annotated for each comparison: (B) Het female vs. WT female; (C) Het male vs. WT male; and (D) Het female vs. Het Male. (E-G) Gene set enrichment analysis (GSEA) is shown for cellular component for:(E) Het female vs. WT female; (F) Het male vs. WT male; and (G) Het female vs. Het Male. (H) Disease enrichment by DisGenNet is shown for DEGs between Het female vs. Het Male comparison. (E-H) Circle color shows adjusted P value and the size of the circles shows the enrichment scores. (I-J) Starbursts plots show disease specific associated DEGs for epileptic encephalopathy (I) and for Severe intellectual disability (J). For both plots DEGs labelled in red are genes with pathogenic variants that confer high-risk for ASD while grey DEGs are neurodevelopmentally important genes.

Next, to further explore ASH1L-driven molecular programs that are male and female specific, we directly compared DEGs from male and female *Ash1L^gt/+^*mice and found 722 DEGs (Figure 7D, Supplement to Figure 7, Supplementary Table S2). GSEA Pathway enrichment analysis showed genes associated with autophagy and protein folding were differentially regulated between sexes in the heterozygous animals. Further GSEA analysis of biological process showed upregulation of sensory function related genes (i.e. sensory perception) and downregulation of genes involved in mitochondrial function (i.e. oxidative phosphorylation) in *Ash1L^gt/+^* females compared to *Ash1L^gt/+^* males (Supplement to Figure 7, Table S3. To define whether the differences identified were solely driven by sex independent of genotype we compared the WT males and females, and identified 599 DEGs (Supplement to Figure 7), Table S2). We found that only one fourth of the DEGs in the *Ash1L^gt/+^*male vs. female comparison overlapped with the WT male vs. female comparison. Further, we found that genes involved in the spliceosomal complex and in translation (i.e. ribosomal subunit) were enriched in both comparisons (Supplement to Figure 8, Table S2). Therefore, while there is a contribution of sex to the molecular signatures observed when comparing male and female *Ash1L^gt/+^)*, the genotype drives a distinct sex-specific transcriptomic signatures in *Ash1L* mutant mice.

**Figure 8:**
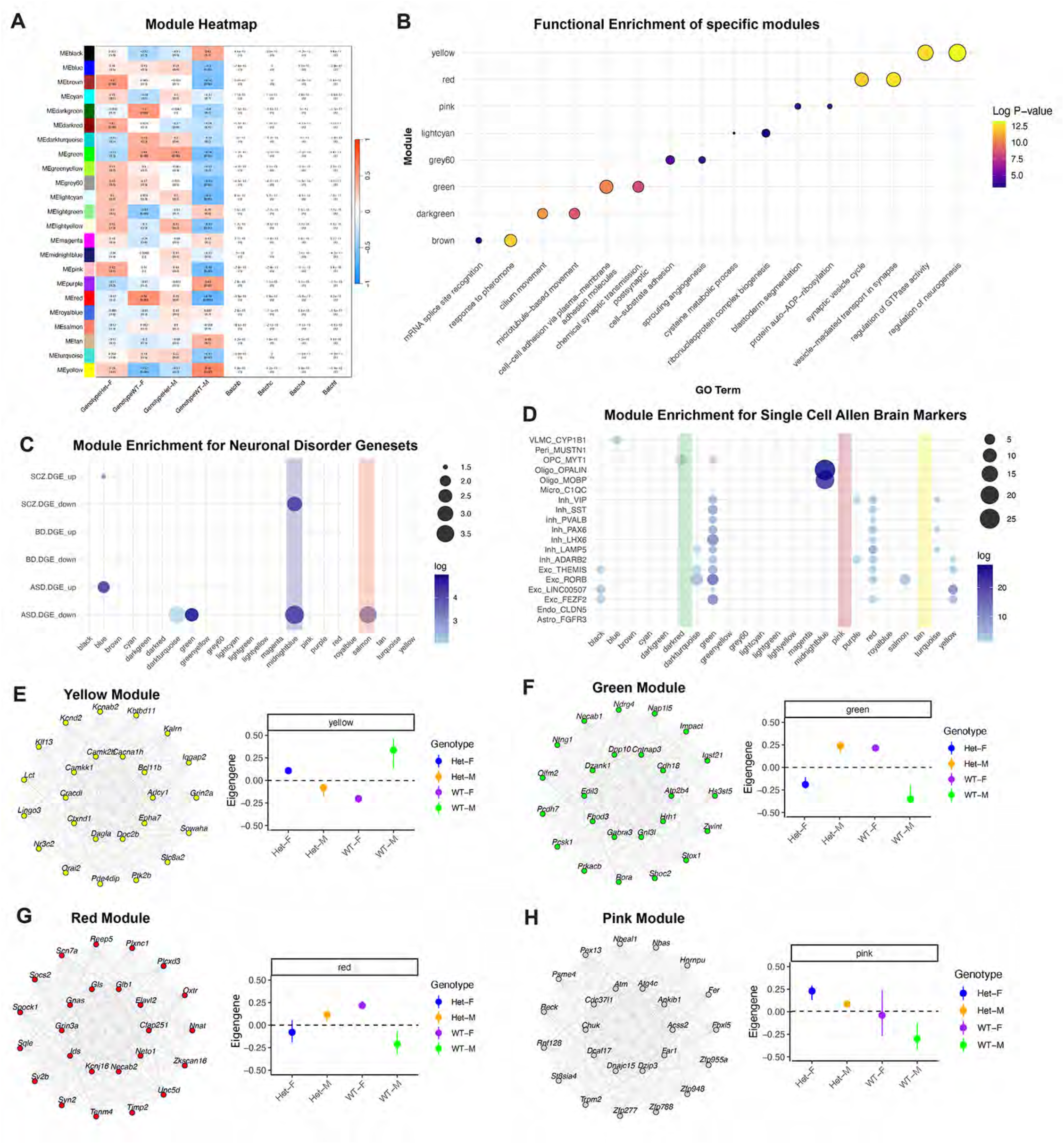
WGCNA shows distinct male and female signatures driven by ASH1L. (A) Heatmap shows changes in expression for each module in relation to genotype and sex. (B) WGCNA functional enrichment analysis for specific modules is shown, with colors depicting adjusted log p-value. (C) WGCNA modules showing disease association with neuropsychiatric disorders highlights upregulated and downregulated DEGs within each module. The size of the solid black circles represents the number of DEGs and the hue color represents the log p-value with darker hues showing greater statistical significance. Solid bars highlight specific modules. (D) Magma plots show WGCNA enrichment for single cell expression from the Allen Brain Atlas. Solid black circles size represent number of DEGs while hue color represents log p-value. Solid bars highlight specific modules. (E-H) Gene networks with relevant hub genes and the directionality of eigengene for sex/genotype are shown for the yellow (E), green (F), red (G) and pink (H) modules.

The identification of distinct molecular programs in males and females in the heterozygous animals led us to examine disease association. We mapped DEGs to their human orthologs and performed enrichment analysis using the DisGeNET pathway database^60^ (Figure 7H, Table S4). The top enriched pathways were associated with epileptic encephalopathy, postnatal microcephaly, autistic behavior, and severe intellectual disability, suggesting distinct susceptibility patterns in female versus male ASH1L mutant mice. Furthermore, the genes driving these pathways were enriched for neurodevelopmental regulators that included genes with pathogenic variant of high-risk for ASD as classified by SFARI^61^ (Figure 7I-J).

### Weighted Gene Co-Expression Network Analysis (WGCNA) reveals sex-specific transcriptional programs driven by ASH1L in relation to developmental stage and brain cell type

To further distil mechanistic insight associated with ASH1L function, we performed weighted gene co-expression network analysis (WGCNA), identifying 23 distinct co-expression modules with eigengene networks associated with each genotype (Figure 8 A-H, Supplement to Figure 8 and Table S5). Functional enrichment analysis revealed module-specific associations with key neurobiological processes, including neurogenesis (yellow module), synaptic vesicle function (red module), postsynaptic transmission (green module), and microtubule-based movement (dark green module) (Figure 8B, 8E-G, and Table S5). Interestingly, sex-specific differences were present across modules, for example, the neurogenesis-associated yellow module was downregulated in *Ash1L^gt/+^* males and upregulated in *Ash1L^gt/+^*females. In contrast, the red and green modules, associated with synaptic function, were downregulated in females and upregulated in males. However, in some modules the directionality of gene expression changes was similar in both males and females (e.g. upregulation in both sexes in the pink and brown modules) (Figure 8H, and Table S4).

To further examine the potential relationship between ASH1L-driven transcriptional changes and complex brain disorders, we compared module-specific gene expression patterns with a neuropsychiatric brain gene expression meta-analysis from the PsychENCODE Consortium^62^. Several modules showed enrichment for genes associated with ASD (green, midnight blue, blue, dark turquoise, and salmon) and schizophrenia (blue and midnight blue) (Figure 8C, Supplement to Figure 8). Similarly, analysis of ASD-associated gene sets showed that the green, light green, magenta, purple, and yellow modules were significantly enriched for ASD risk genes (Supplement to Figure 8). Finally, as single cell studies on an ASH1L mutant mice have shown cell-type specific signatures^51^, we analysed single-cell datasets from the Allen Brain Atlas to dissect potential contributions of different cell types to the phenotypes observed in *Ash1L^gt/+^.* We found enrichment of ASH1L-regulated genes in excitatory and inhibitory neurons (black, dark turquoise, green, red, and yellow modules), as well as in oligodendrocyte precursor cells (OPCs) and microglia (dark red and midnight blue modules) (Figure 8D). Taken together these findings suggest that ASH1L-mediated transcriptional regulation affects key neurodevelopmental pathways in a sex-dependent manner, cell type specific manner, and may contribute to the genetic risk architecture of ASD and schizophrenia.

## Discussion

This study investigated the role of ASH1L mutations in a neuro-developmental disorder. Utilizing both C57BL/6 inbred and CD-1 outbred mouse models, we found that these mutations lead to significant brain phenotypes, including epilepsy, microcephaly, and altered dendritic morphology. Notably, the study highlights the influence of genetic background on the effects of ASH1L, revealing that different mouse strains exhibited varied responses to the mutation. Furthermore, the analysis of human subjects indicated sex differences in the manifestation of epilepsy and autism, with epilepsy being more prevalent in females and autism in males. This suggests that the neurodevelopmental impacts of ASH1L mutations are not only influenced by genetic background but are also dependent on biological sex. Electrophysiology revealed distinct differences in ASH1L effects on hippocampal neuron function between male and female mice, indicating that the circuit-level effects of ASH1L mutations may be modulated by biological sex. In summary, this study underscores the complexity of ASH1L-associated neurodevelopmental disorders, particularly the importance of considering both genetic background and sex.

### Effect of genetic background causes the most variability in inhibitory systems

The effect of genetic background is frequently cited as the cause of variability in disease penetrance. We chose the standard C57BL/6 mouse that is used in neuroscience studies specifically with regard to behavior. We also performed most studies in the CD-1 background mouse, which is an outbred strain where we would expect a larger degree of variability. Histone modifications, as regulated by ASH1L, can alter the expression of many genes, so we expected that these different genetic backgrounds would modulate the degree of the appearance of certain phenotypes. This was true to a certain degree. For example, the dendritic branching phenotype is more severe in the CD-1 background, where disruption of ASH1L resulted in simplification of both apical and basal dendrites at later time points, but only in basal dendrites in the B6 mice. Background can completely suppress certain phenotypes. While children with NDD have a high rate of sleep disorder, only mice with ASH1L mutations on the CD-1 background had fragmented sleep. Interestingly, the effects of genetic backgrounds appear to be much more variable on inhibitory neurons, with opposing effects on some systems. Numbers of somatostatin cells are decreased in the BL6 background, while increases are observed in CD-1 brain regions. Disruption of ASH1L alters PV inhibitory synapses in different directions depending on background. This degree of variability in inhibitory neurons was surprising and may suggest that in some cell types genetic background confers selective vulnerability in disease states.

### Importance and limitations of human studies and animal modeling in neurological and developmental disorders

In understanding ASH1L-associated NDD, we analyzed both rodent models and human subjects. One limitation of the rodent studies was the lack of behavioral phenotypes, where in the cohort of human subjects, we were able to assess patients with respect to intellectual disability and autism, and perform a neurological examination with good fidelity. In previous behavioral studies of ASH1L, phenotypes were observed after conditional deletion of both copies of ASH1L in only the central nervous system. While this work was important for understanding the role of ASH1L, full loss of function is not observed in humans. It is likely that human disorders with incomplete penetrance of neuropsychiatric symptoms or intellectual disability falling in the mild/moderate range may be difficult to model in rodents using standard behavioral studies.

Despite the lack of behavioral phenotypes in the rodent system, the model remains important for understanding the fundamental biology of ASH1L in the developing brain. The availability of well-characterized mouse strains allowed us to perform studies looking at the effects of genetic backgrounds in a controlled manner across both structural and functional domains. In addition, we were able to discern developmental effects of ASH1L at a cellular level *in vivo*, which also validated findings in human iPSC-derived neurons. Finally, the cerebellar changes in mouse models across both backgrounds led us to perform neurological assessments that focused on fine coordination and balance, which was impaired in a high proportion of human subjects. Taken together, human studies along with animal studies provide greatest depth and breadth in discerning the roles of ASH1L in health and disease.

### Sexual Dimorphism in neurodevelopmental disorders

Sexual dimorphism in neurodevelopmental disorders is well-described, especially with regard to autism, intellectual disability, and epilepsy. In general, boys are more severely affected in these disorders. Overall, this increased severity appears to be true in ASH1L-related disorders, as well. Autism was more prevalent and more severe in boys, and developmental delays were also more severe in boys than in the girls, especially in the motor domain. Furthermore, the parental rating scales for adaptive function suggest that overall, boys have more difficulty, reflecting the consensus view that boys are more severely affected in neural developmental disorders. While male susceptibility is often reported for X-linked disorders, the ASH1L gene is autosomal, thus, sex differences must occur via other mechanisms.

In contrast to the autism phenotype/finding, the epilepsy phenotype is selectively more prevalent and more severe in girls. In the literature, epilepsy is reported in up to 20% rate in people with autism^63^. In ASH1L-affected boys in our study, five of ten boys had epilepsy, a history of seizures or abnormal EEG in our study. However, ***none*** of the boys have epileptiform activity by EEG at baseline, and only one boy has epileptiform activity by hyperventilation on EEG. In contrast, *eight of thirteen girls have epileptiform activity by EEG* and nine of the thirteen girls have either a history of epilepsy, current epilepsy, or epileptiform activity on a standard baseline clinical EEG. While people can have epilepsy in absence of abnormal EEGs, in general the presence of epileptiform features by EEG is a predictor of the severity of the epilepsy^64^. It is remarkable that the sex difference in epilepsy was not previously recognized, but that is likely due to the fact that the female phenotype was largely unrecognized. Of the 14 case reports for which there is detailed clinical information, only two include girls^8^.

The clinical presentation of our patient cohort as well as electrophysiological hippocampal signatures in Ash1l heterozygous animals are sex-specific and correlate with distinct transcriptomic signatures in male and female mice that are important for the development of neural circuits. Mitochondrial biogenesis and function have been suggested to be differentially regulated in male and female brains^65^. Our transcriptome studies suggest that Ash1l could be driving the differential regulation of mitochondrial function in a sex-specific manner. Additionally, we find that Ash1l differentially regulates genes involved in either autophagy and proteasome function in females vs. males. Previous work has shown sex-specific differences in autophagy are associated with Alzheimer’s disease as well as cardiovascular disease^66^. While cardiovascular phenotypes were previously reported in a subset of ASH1L patients^9^, there is currently no clinical evidence to support an association of Alzheimer’s disease with Ash1L. However, as the patient population ages, longitudinal studies may define potential sex-specific neurodegenerative presentations in relation to ASH1L disease.

### How does ASH1L cause a sex-specific circuit defect

The sex-specific hyperexcitability of hippocampal neurons in female mice likely arises from similar mechanisms causing the increased epilepsy in girls. How ASH1L causes a sex-specific circuit defect is at present unknown, however, this effect could be cell autonomous/ non-autonomous/ or both. ASH1L may affect circuitry directly in female mice via altered gene expression. ASH1L counteracts PRC2 methylation of H3K27me3. When there is less ASH1L activity, PRC2 activity is dysregulated. Since PRC2 binds XIST, the non-coding RNA which is the main repressor of gene expression for X-inactivation, it is plausible that ASH1L may indirectly alter XIST activity in neurons from female subjects. Alternatively, ASH1L disruption may alter sex-specific signaling in brain, by either altering the response to sex-specific steroids or by impairing the production of sex-specific signaling molecules themselves. The underlying causes for these differences are an area of active investigation, and electrophysiology in mice will allow us to investigate the sex difference in a tractable model system.

## Supporting information

RNAsequencing

Human subjects data

## Acknowledgments

We would like to thank the patients and families who participated in this study and especially the family organization, CARE4ASH1L. We also acknowledged the special efforts of predoctoral students who helped us in the laboratory:. We thank our funding agencies for supporting this study. This work was supported with grants from the Autism Science Foundation, Eagles Autism Foundation Grant, NIH/NIMH:1R01MH127081-01A1, and NIH/NIMH:1R21MH136643-01.

## Author contributions

CP, SBL, and JSL conceived of and designed the study, wrote the manuscript. SBL and JSL led the funding acquisition. CP and EN led the mouse studies. EN, JE, CP performed the collection and the analysis of the clinical data. EN, LG performed mouse EEG studies. DN performed the analysis of epilepsy phenotype and EEG. JSL performed the neurological examinations. BK performed the neuropsychological assessments of the patients. HH performed the whole cell patch clamp electrophysiology. MJ performed RNA isolation, library preparation data analysis, and manuscript writing. SD assisted with library preparation.

## Materials and Methods

All clinical activities were carried out in strict accordance with the U.S. Department of Health and Human Services Office for Human Research Protections regulations and were approved by the Lifespan Hospital Institutional Review Board (protocol 1680205). Participants were enrolled prospectively in our study based on self-referral by families of the participants. All participants were confirmed to have a pathogenic or likely pathogenic variant of ASH1L based on clinical genetic testing and confirmation by research team. Informed consent was obtained from parents or guardians of pediatric participants in the study, and assent was obtained from pediatric participants when appropriate. During site visits, participants and families underwent a structured interview with research staff, behavioral and cognitive assessments, a neurological exam and a standard clinical EEG. Participants also signed a medical release form, and additional information was obtained from medical records including clinical genetic testing, neuropsychological evaluations, speech and language evaluations, individual educational plans, evaluations by specialists including neurology, endocrinology, gastroenterology, cardiology, urology, and others, imaging studies, and laboratory studies. All data was maintained in REDCap and the Lifespan Secure Server using de-identified participant codes.

The **Autism Diagnostic Observation Schedule**, Second Edition (ADOS-2) is a standardized interview and observation of ASD symptoms, including repetitive restricted behaviors and social affect/communication. All ADOS-2 administrations were conducted by assessors who were trained to research reliability. The specific ADOS-2 module was selected based on the age/abilities of the given participant. The autism/ASD cut-off criteria were calculated as described in the ADOS-2, and this dichotomous variable was utilized in this study to reflect a positive or negative ADOS finding and/or whether the ADOS confirmed or disconfirmed a clinical diagnosis. To measure ASD severity, the comparison score was utilized. The comparison score ranges from 1-10, with 1 indicating minimal-to-no evidence of autism-related symptoms and 10 indicating a high level of impairment.

The **Wechsler abbreviated scales of intelligence, second** edition (WASI-II; Wechsler 2011), four -subtest version, was utilized for all 6+ year old participants. The Wechsler primary preschool scales of intelligence, fourth edition (WPPSI-IV) was administered for younger participants (in this study, 4-5 year olds). The overall intelligence, verbal intelligence, and visuospatial intelligence composite scores were utilized to assess intelligence.

The **Social Responsiveness Scal**e, second edition(SRS-2) is a caregiver-completed survey designed to quantify symptoms typically associated with Autism Spectrum Disorder (ASD). Three versions of the SRS-2 exist and are distributed based on the patient’s age: SRS-Preschool, SRS-School Age, and SRS-Adult, but all versions contain the same material and calculations. If a participant does not fit within the age categories described above, an SRS-2 is not given for evaluation. While there are sex specific SRS-Adult forms, these are not utilized in this study.

There are 65 questions total with each question belonging to symptom subcategory: Awareness, Cognition, Communication, Motivation, and Restricted Interest and Repetitive Behavior (RRB). Each question is scaled from 1-4 (1= not true, 4 = almost always true) and can only have one answer per question.

Raw scores are added together to calculate the SRS Total Score. The Social Communication and Interaction (SCI) score is calculated by subtracting the RRB raw score from the Total T-score. Raw scores are then converted to age matched “T-scores” using the SRS-2 published grading manual (source). Only the T-scores from SCI, RRB, and Total T-scores are used in determining ASD DSM-5 criteria.

The **Vineland Adaptive Behavior Scales,** Third Edition, assesses an individual’s adapted behaviors due to an intellectual and/or developmental disabilities. All ages (0-99) can be evaluated, but only those ages 9 and under will have their motor skills scored. Multiple versions of the Vineland-3 are available for use, including self-reported measures, but the parent/caregiver comprehensive form is utilized in this study. This survey aims to evaluate an individual’s ability to communicate (verbally and through reading and writing), socialize (both in groups and with themselves), their daily living skills (hygiene and self-sufficiency), and motor ability (both gross and fine motor skills).

Each aim (communication, daily living, social skills, and motor skills) has three subcategories that are graded based on the chosen answer: “Usually or Often” = 2 points, “Sometimes” = 1 point, or “Never” = 0 points. Additionally, there are some yes/no questions in each subcategory where “yes” = 2 points, and “no” = 0 points. Only one answer is allowed per question.

Raw scores are calculated through adding up each subcategory, which are then translated into V-scale scores through the published Vineland-3 grading manual [Cicchetti, Saulnier, Sparrow, NCS Pearson Inc, 2016]. V-scale scores of each subcategory are added together to create the cumulative “standard” scores and correlative percentile ranks. Finally, an “Adaptive Behavior Composite” is calculated through adding up the standard scores of the communication, daily living, and social skills categories.

V-scores in each category above a score of 86 are considered adequate, as the “average” is 100 with a standard deviation of +/-15. A score 85 and below is considered moderately low, with scores below 70 being considered especially low [Health Net Federal Services (HNFS) LLC/Tricare, Vineland-3].

The **Social Communication Questionnaire** (SCQ) aims to evaluate a participant’s social communication skills as observed by a primary caregiver. As social skill deficiencies are common among those diagnosed with autism spectrum disorder (ASD), the SCQ can be used as an investigative tool evaluating a patient’s autism severity. The SCQ is separated into two categories of questions: one referring to a participant’s entire lifetime and one referring to only the last 12 months, with each category having 20 questions. The SCQ questions do not change between age groups, but it is not given to primary caregivers with participants <4 years old.

The **Aberrant Behavior Checklist** (ABC) is a caregiver-completed, investigative tool used to evaluate nontypical behaviors in individuals above the age of 6 with intellectual and developmental disabilities. Multiple versions of this survey are available, but the 31-item version, focusing on irritability and hyperactivity, was utilized for this study. The checklist offers 4 choices per question, with only one answer per question, and is graded on a scale from 0 points (not at all) to 3 points (severe). Questions 1-15 add up to make the irritability sub-scale score and questions 16-31 add up to make the hyperactivity sub-scale score with each sub-scale having up to 45 total points. The larger the score, the more severe symptoms are exhibited, as seen by the primary caregiver.

### Electroencephalogram (EEG) recording in human patients

Participants received EEG studies in the Rhode Island Hospital pediatric EEG laboratory. 21-lead EEG electrodes were placed according to the international 10-20 system of electrode placement. Digital recording was obtained at 256 Hz sampling rate during the awake and sleep states, with time-locked video and single lead electrocardiogram (ECG). For visualization by anterior-posterior bipolar and average montages, low-pass filter is set at 70 Hz and high-pass filter at 1 Hz, sensitivity 30 uV/mm, time base 15 mm/sec. Three minutes of adequate hyperventilation was performed when age-appropriate in absence of contraindications, according to standard protocol. Photic stimulation using a stepwise increase in photic frequency between 2 and 30 Hz was performed when appropriate.

### Mouse models

Animals were housed in an Association for Assessment and Accreditation of Laboratory Animal Care (AAALAC) accredited facility. All animal care and procedures were carried out in strict accordance with National Institutes of Health Guide for the Care and Use of Laboratory Animals and were approved by the Institutional Animal Care and Use Committee at Brown University (protocol 20-10-002). ASH1L^gt/+^ mice were obtained from The Jackson Laboratory, strain #028220. Founder mice were backcrossed for three or more generations onto either a C57BL/6J (The Jackson Laboratory, strain #000664) or CD-1 (Charles River Laboratories, strain #022) background.

The genotype of experimental mice was determined with polymerase chain reaction (PCR). A common for-ward primer (5’-GAGAGACGGCTCAGTGTTGA-3’) was paired with reverse primers to amplify the WT allele (5’-TGTTGGATTTAGTGTGTGTGTGTG-3’) or the gene trap allele (5’-ACCTGGTTGTCATGGAGGAG-3’). The PCR conditions were 95C for 3 min followed by 34 cycles of 95C for 30 sec, 62C for 45 sec, and 72C for 1 min, followed by a final extension of 72C for 5 min. PCR products were visualized on a 1.5% agarose gel. The WT primer set amplified a 191 bp fragment, and the gene trap primer set amplified a 223 bp fragment. Once during the experiment, the mutant PCR product was also confirmed by Sanger sequencing.

### Histology and immunohistochemistry

For mouse tissue expression studies, at least 4 separate mice per genotype were used for each reported result. Mice at P0 were anesthetized with ice and intracardially perfused with 0.1 M PBS (pH 7.4) followed by 4% paraformaldehyde (wt/vol). Brains were removed and postfixed in 4% paraformaldehyde overnight, weighed, and photographed. Brains were then embedded in paraffin and sectioned into coronal sections (5 μm) on a Leica RM2125 RTS microtome. Before staining, sections were deparaffinized and antigen retrieval was performed by incubating slides at 100C for 45 minutes in a sodium citrate buffer (10 mM sodium citrate, 0.05% Tween-20, pH 6.0). Sections were then briefly washed in PBS before undergoing the staining procedure described below.

Mice at P14 and 2 months were anesthetized with sodium pentobarbital and intracardially perfused with 0.1 M PBS (pH 7.4) followed by 4% paraformaldehyde (wt/vol). Brains were removed and postfixed in 4% paraformaldehyde overnight, weighed, and photographed. Brains were then transferred to 30% sucrose solution until saturated for cryoprotection. Coronal tissue sections (40 μm) were sectioned using a microtome (Thermo Scientific Sliding Microtome Microm HM 430) and collected in PBS. Immunocytochemistry at P14 and 2 months was performed on free-floating sections. All sections were incubated for 1 hour in blocking buffer (PBS containing 10% normal goat serum (vol/vol) and 0.5% Triton X-100 (vol/vol)). Sections were incubated with agitation at 4C overnight in primary antibodies diluted in 0.1 M PBS (pH 7.4) containing 5% normal goat serum (vol/vol) and 0.5% Triton X-100 (vol/vol)). See supplemental information for a list of the primary antibodies used in this study. Secondary antibodies were selected according to the species of the primary antibodies used, and used at a 1 to 500 dilution. Sections were incubated with secondary antibodies for 2 hours at room temperature followed by DAPI for 10 minutes and mounted on slides using Fluoromount-G (Cat#0100-01, Southern Biotech, Birmingham, AL). All brightfield imaging was performed with a Nikon Ti2E Inverted Microscope. All fluorescence imaging was performed with an Olympus FV3000 confocal microscope. Images were processed using Photoshop and ImageJ (NIH).

### Image analysis

All image analysis was either performed by an independent, blinded observer, or performed in a blinded manner by using the RandomNames plugin. For all image analysis, at least 2-3 sections per mouse and 3-4 mice per genotype were used. For cell counts in the cortex, somatosensory cortex was imaged at 20x. Images were cropped into sections of equal width (100 μm at P0 and 500 μm at P14 and 2 months). Images were then divided into 10 equal regions of the cortex spanning from the pial surface to the ventricular zone of the cortex. The number and percentage of cells in each box were counted and compared. The area of each box was measured to calculate cell density. For cell counts in the hippocampus, the hippocampus was imaged at 20x bilaterally. Hippocampal landmarks were identified as described in Whitebirch et al. 2022 ^67^.

For perisomatic synapses, layer V cortex was imaged at 60x. Two fields of view per section, two sections per mouse, and four mice per genotype were used. A single z plane was used for analysis. Per field of view, twelve cells in the center of the field of view were randomly selected and provided to an independent observer. In total, about 384 cells per genotype were analyzed.

### In-situ hybridization

In-situ hybridization was performed on free-floating sections from P14 mice prepared as described above. Staining was performed according to manufacturer’s instructions using the ACDBio RNAscope Intro Pack for Multiplex Fluorescent Reagent Kit v2-Mm (catalog #323136) and probes against ASH1L (#567141), TUBB3 (#423391-C3), and ALDH1L1 (#405891-C2). For visualizing mRNA, Opal fluorophores were purchased from Akoya Biosciences (Opal 520, PN FP1487001KT; Opal 570, PN FP1488001KT; Opal 690, PN FP1497001KT).

### Golgi analysis

2-month-old mice were perfused intracardially with a normal saline solution and brains were dissected. Golgi staining was performed with the FD Neurotechnologies FD Rapid GolgiStainTM Kit (PK401) according to manufacturer instructions^68^. Brains were sectioned to a thickness of 100 μm using a Leica CM1860 cryostat. Layer V pyramidal cells were imaged at 20x magnification with a Nikon Ti2E Inverted Microscope. Neurons were traced manually in three dimensions in Photoshop. Neurite length, number of branch points, complexity index, soma area, and Sholl analysis were analyzed by an independent observer using Neurolucida software.

### Whole cell patch clamp electrophysiology in CA1 hippocampus and Layer 5 somatosensory cortex in mice

Whole cell patch clamp electrophysiology was performed on male and female mice between postnatal days 23-26. Studies were performed in daylight hours between noon and 4pm. Coronal sections of 300μm in thickness were prepared in an ice-cold and oxygenated high-sucrose slicing solution and were then transferred to oxygen-saturated ACSF containing the following (in mM): 126 NaCl, 26 NaHCO3, 10 glucose, 2.5 KCl, 1.25 NaH2PO4.H2O, 2 MgCl2.6H2O, and CaCl2.2H2O; pH 7.4. All sections were incubated at 32°C for one hour prior to recording at room temperature. We followed previously described methods for obtaining electrophysiological parameters for active and passive membrane properties including input resistance, resting membrane potential, capacitance, rheobase, action potential threshold, amplitude and frequency^75^. All recordings were made from CA1 of the hippocampus or Layer V of somatosensory cortex. A high chloride intracellular solution was used to study the intrinsic properties of neurons (in mM): 70 Kgluconate, 70 KCl, 2 NaCl, 10 HEPES, 4 EGTA, 2 Na2-ATP, 0.5 Na2-GTP. Recordings were performed using (Multiclamp 700B, Molecular Devices), and digitized (DigiData 1550B, Molecular Devices). Measurements of intrinsic properties were analyzed using Clampfit software (v. 11.2 Molecular Devices). We used the python-based open-source tool Google Colaboratory to generate and analyze action potential (AP) phase plots from WT and Ash1L^+/gt^ animal electrophysiological recordings. Axon Binary Format (ABF) files were read and analyzed by the Python 3.10 and pyABF package (ref 1). We selected the 80pF current injection point for AP phase plot analysis.

### Electroencephalography depth electrode implantation and recording

Adult female and male mice (2 months old) were anesthetized with a constant stream of isofluorane. Once the mouse was unresponsive to toe pinch cues, the head was shaved and a moistening agent was applied to the eyes to prevent the eyes from drying out. The shaved surgical site was thoroughly sterilized, and lidocaine was injected subcutaneously at the surgical site. The skull was then exposed and chelated with 3% hydrogen peroxide. iBond was applied to the surface of the skull and hardened with UV light exposure. Three holes were then drilled into the skull: one above the right hippocampus (1.5 mm posterior to bregma and 1.5 mm from the midline), one above the left frontal cortex (1.5 mm posterior to bregma and 1.5 mm from the midline), and one above the cerebellum (1 mm from lambda at the midline). A headset attached to 3 insulated electroencephalography (EEG) wires and 2 electromyography (EMG) wires was then lowered stereotactically, such that a wire lead was placed into each of the 3 holes, and the EMG leads are put into contact with the muscle tissue of the back. The holes, surface of the skull, and region immediately surrounding the surgical site were covered up with dental cement. The animals were allowed to recover for 5 to 7 days after surgery. Behavioral, EEG, and EMG observations were made by using the Pinnacle 8400 three channel monitoring system with simultaneous video recording (Pinnacle Technology). Mice were subjected to at least 72 hours of continuous recording. Recordings were sampled at 2kHz and low-pass filtering was done at 100Hz. EEG signals were collected and analyzed with Sirenia software. Interictal spikes were defined as single spikes with an amplitude of 2x baseline in the absence of EMG activity and without visible movement by the mouse immediately preceding the spike. To avoid artifacts from movement of the mouse, interictal spikes were only counted when the mouse was sleeping. Power spectral analysis was performed with Sirenia Seizure software.

### Sleep analysis

was scored manually from 7am to 7pm on the second day of recording using Sirenia Sleep software. Bin size was 10 seconds. If the mouse entered a sleep or wake state for less than 30 seconds total, that was disregarded to avoid artificially decreasing bout length.

### Behavior

All mice are housed in a barrier facility with a 12-hour light/dark cycle and ad libitum access to food and water. Except as described, mice were tested at 6-8 weeks of age. Mice were handled for 10 minutes on three consecutive days prior to the start of behavioral testing. Except as described, all behavioral experiments were performed between 2-4PM (ZT 7-9).

### Open field

Mice were placed in a 40 cm x 40 cm x 30 cm arena and allowed to habituate for 10 minutes. Mice were then recorded exploring the arena for an additional 10 minutes. Videos were analyzed using DeepLabCut, a free toolbox for pose estimation from the Mathis laboratory ^69^. The snout, left ear, right ear, and base of tail were labeled as coordinates of interest. Twenty randomized images per experiment from 8 separate experiments were used as a training dataset. The model was trained for 200,000 iterations with a final loss of 0.0018. Distance traveled and percent time in center were calculated from the resulting snout coordinates using Python (code available upon request).

### Novel object recognition

At 7PM on day 1, mice were habituated to the area for ten minutes. At 7AM the next day, two identical objects (either a tower made of Legos or a T25 tissue culture flask filled with sand) were placed in the arena, and the mouse was allowed to explore the objects for ten minutes. 12 hours later, at 7PM on day 2, one object was replaced with a novel object and the mouse was placed in the arena for ten minutes. Objects were randomized and counterbalanced, and all objects were cleaned and sterilized after every stage of every trial. All stages were video recorded. Interaction time was measured with a stopwatch by a blinded, independent observer. Preference index was calculated as (time interacting with novel object) / (total time interacting with both objects)^70^.

### Social interaction test

Same-sex, weight-matched (± 5 grams) non-littermate unfamiliar mice were used as the probe mouse and novel mouse. Each test mouse was allowed to habituate in the center chamber of a three-chamber testing apparatus for 10 minutes. Probe mice were also habituated to the plastic cage for 2 x 15 minutes and screened for aggressive behaviors. At the end of the habituation period, test mice were allowed to explore all three chambers of the apparatus for 10 minutes. After the habituation period, the test mouse was guided back to the center chamber and the probe mouse was placed in one of the two sides. An empty plastic cage was used as the object and placed into the other side. The side of the chamber containing the probe mouse was randomized and counterbalanced. The test mouse was then allowed to explore all 3 chambers for 10 minutes. At the end of the first stage, the probe mouse and test mouse were each returned to a clean, empty cage and allowed to rest for 10 minutes. For the social novelty test, the probe mouse was returned to the plastic cage, as well as a novel mouse in the second plastic cage. The test mouse was then allowed to explore all 3 chambers for 10 minutes. All phases were video recorded. Interaction time was measured with a stopwatch by a blinded, independent observer. For social preference test, social preference was calculated as (time interacting with mouse)/(total time interacting with object and mouse) ^70^.

For social novelty test, novelty preference index was calculated as (time interacting with novel mouse)/(total time interacting with both mice).

### Grooming

Mice were placed in a clear chamber with 1cm of fresh bedding and allowed to habituate for 10 minutes. Mice were then recorded for an additional 10 minutes. Grooming time was measured with a stopwatch by a blinded, independent observer ^71^.

### Marble burying

A clean, empty cage was filled with 5 cm of fresh bedding. Twenty glass marbles were evenly spaced in 4 rows. Each mouse was allowed to bury the marbles for 30 minutes, after which the mouse was returned to its home cage. Number of marbles buried was scored by three blinded, independent observers, and the results were averaged. A marble was considered “buried” if more than two-thirds of the marble was covered by bedding ^72^.

### Righting reflex

Pups were placed in a supine position on a flat, clean surface. The time to return to a prone position was measured with a stopwatch. A pup failed the test if its time to right exceeded 1 minute. Mice were tested at postnatal day 4, 7, and 10^45^. Each pup received two trials at each postnatal time point and allowed to rest in its home cage for 2 minutes between trials. The two trials were averaged to calculate the righting time.

### Hippocampal tissue collection, Library Preparation and Sequencing

At day P26. Hippocampi were dissected and collected in RNAase/DNAase free microcentrifuge tubes and flash frozen in liquid nitrogen and stored at -80°C until all samples from all genotypes and sexes were collected. Hippocampi collection followed a strict circadian protocol by collecting samples between 9 and 11 AM to reduce variability in the transcriptome due to time of the day collection of samples. RNA was extracted from tissue samples using the RNeasy Micro Kit (Qiagen, #74004) following the manufacturer’s protocol, with mechanical lysis performed by trituration using a needle and syringe. RNA quality was assessed for each sample using the Agilent 2100 Bioanalyzer, and only samples with RNA Integrity Numbers (RIN) >9.5 were used for downstream analysis. Stranded cDNA libraries were prepared using the Collibri™ Stranded RNA Library Prep Kit for Illumina™ Systems (Invitrogen, #A38996096). Sequencing was carried out on an Illumina NovaSeq platform with 150-bp paired-end reads, achieving a depth of 100 million reads per sample.

### RNA-Seq Data Processing and Analysis

Raw sequencing data quality was evaluated using FastQC (version 0.11.9) and TrimGalore (version 0.6.6). Reads were aligned to the Gencode reference genome M36 (GRCm39) using Salmon (version 1.5.2) in the mapping-based mode for transcript-level quantification. Count matrices were generated from transcript abundance estimates using the tximport package (version 1.30.2). The RNA-Seq dataset has been deposited in the Gene Expression Omnibus (GEO) under accession number GSE#.

Differentially expressed genes (DEGs) were identified using DESeq2 (version 1.32.0), with statistical significance defined as an adjusted p-value < 0.05 and an absolute fold change ≥1.5. Pathway enrichment analyses, including Gene Set Enrichment Analysis (GSEA) and DisGeNET, were performed using ClusterProfiler (version 4.10.1) and DOSE (version 3.28.2) in R. For disease-associated gene enrichment, mouse gene lists were converted to human orthologs using the msigdbr package (version 7.5.1). Autism spectrum disorder (ASD)-associated genes were obtained from the SFARI database.

### Co-expression Network Analysis

To identify modules of co-expressed genes, we performed weighted gene co-expression network analysis (WGCNA, version 1.73). A soft-thresholding power was automatically selected to achieve a scale-free topology (scale-free topology fit index R² >0.85). Networks were constructed using the blockwiseConsensusModules function with biweight midcorrelation (bicor). Modules were detected using the dynamic tree-cutting algorithm with a deep split parameter of 4 to enhance module specificity. We generated 200 permuted networks to assess network robustness and compared gene connectivity between observed and randomized datasets. None of the permuted networks recapitulated the connectivity patterns of the actual network, supporting the biological relevance of the identified modules. Module eigengenes were correlated with phenotypic traits using Spearman’s rank correlation.

### Functional Enrichment

Functional annotation of module genes was performed using GOstats (version 2.56.0) and Gene Ontology (GO) databases, with all expressed genes (n = 13,412) serving as the background. The overrepresentation of GO terms was tested using a one-sided hypergeometric test, and p-values were adjusted for multiple comparisons using the Benjamini-Hochberg method.

### Neuropsychiatric Gene Analysis

Module gene lists were converted to human orthologs using the msigdbr package (version 7.3.1) for comparison with neuropsychiatric disorder-associated gene sets. ASD-associated genes (SFARI categories 1–3), as well as gene sets linked to schizophrenia (SCZ) and bipolar disorder (BD), were obtained from publicly available resources.

### Gene-Set Enrichment Analysis for cell-type specificity

Cell-type-specific enrichment was assessed using marker genes from the Allen Brain Atlas. Fisher’s exact test (alternative = “greater”, confidence level = 0.95) was applied to evaluate overrepresentation, with results reported as odds ratios (OR) and Benjamini-Hochberg adjusted p-values (FDR).

---

**Figure S1:**
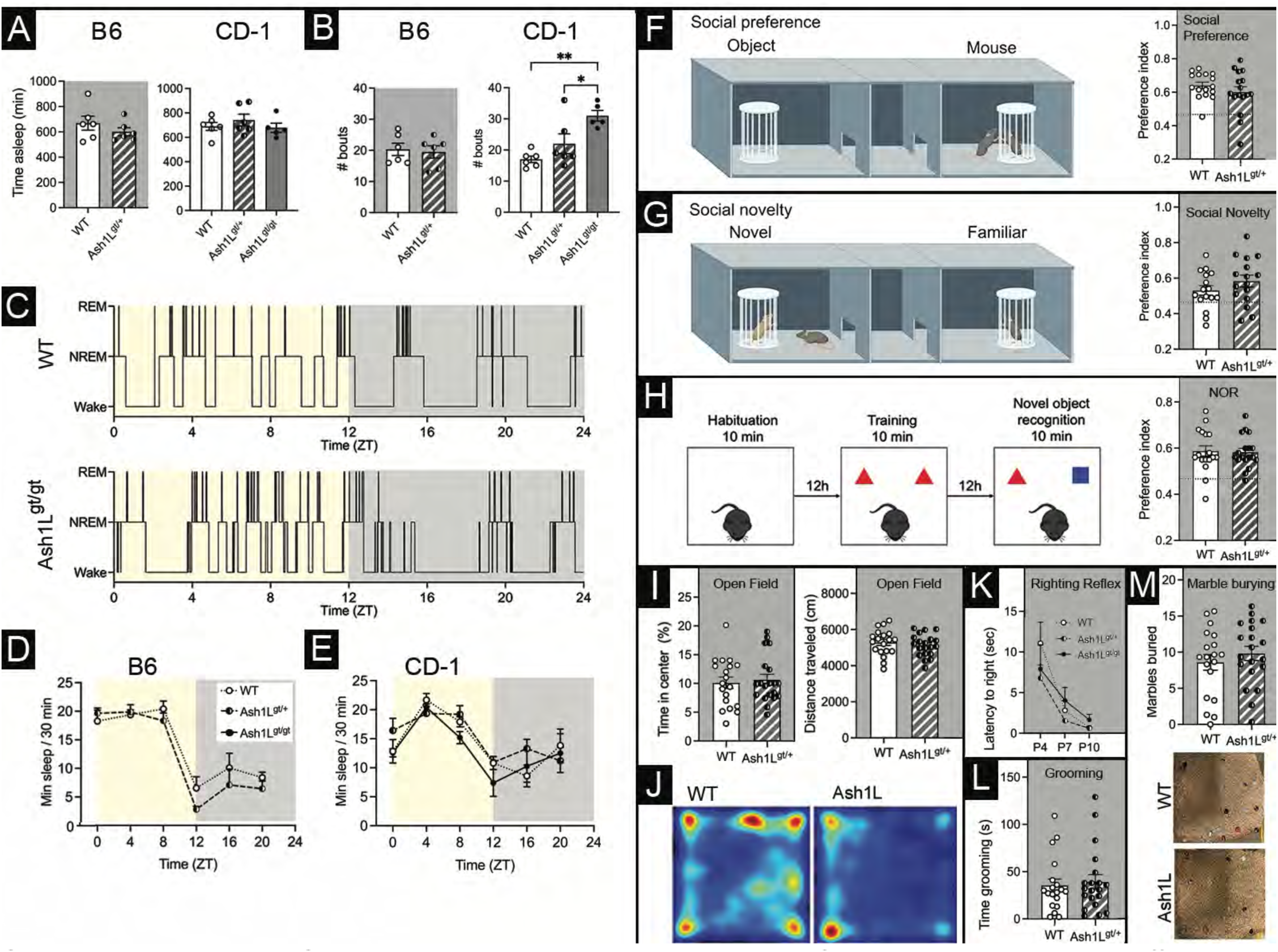
Sleep and negative behavioral data: ASH1L mutant mice have differences in sleep behavior. (A) On a B6 background, ASH1L mutant mice do not have significant differences in the number of sleep bouts or total sleep duration. (B) On a CD-1 background, ASH1L mutant mice sleep the same amount in total duration, but spread across almost twice as many sleep bouts as WT mice. (C) Representative hypnograms of a WT (top) and *Ash1L^gt/gt^* (bottom) mouse showing transitions between wake, non-rapid eye movement (NREM) sleep, and rapid eye movement (REM) sleep during light (left, yellow) and dark (right, gray) phases. (D-E) ASH1L mutant mice on both the B6 (D) and CD-1 (E) backgrounds do not have differences in circadian rhythm. Minutes of sleep per 30 minutes of recording are shown during the light (yellow, left) and dark phase (gray, right). (F) Example of social preference paradigm. Test mouse may interact with a peer mouse (right) or an object (left). ASH1L mutant mice have no significant difference in preference for social interaction. Data analyzed by unpaired t-test (t_30_ = 1.066, p = 0.2950). N = 16 per genotype. (G) Example of social novelty paradigm. Test mouse may interact with a novel mouse (left) or a familiar mouse (right). ASH1L mutant mice have no significant difference in preference for social novelty. Data analyzed by unpaired t-test (t_30_ = 1.235, p = 0.2262). N = 16 per genotype. (H) Example of the novel object recognition (NOR) paradigm. Mice are allowed to habituate to the arena for 10 minutes on the day prior to testing. On the day of testing, mice explore an arena with two identical objects for 10 minutes. 12 hours later, the mouse is allowed to explore an arena with the familiar object and the novel object. A mouse with intact memory is expected to spend more time interacting with the novel object. ASH1L mutant mice have no significant difference in novel object recognition. Data analyzed by unpaired t-test (t_33_ = 0.1730, p = 0.8637). N = 17-18 per genotype. (I) ASH1L mutant mice have no significant difference in percent time spent in the center of the arena during the open field test. Data analyzed by unpaired t-test (t_37_ = 0.3992, p = 0.6921). N = 19-20 per genotype. ASH1L mutant mice have no significant difference in distance traveled during the open field test. Data analyzed by unpaired t-test (t_37_ = 0.7697, p = 0.4464). N = 19-20 per genotype. (J) Representative heat maps showing travel during the open field test by WT (left) and *Ash1L^gt/+^* (right) mice. (K) ASH1L mutant mice have no significant difference in latency to right in the righting reflex test at postnatal day 4 (P4), postnatal day 7 (P7), and postnatal day 10 (P10). Mixed effects analysis (age, genotype) shows no effect of genotype (F_1.604,142.8_ = 1.508, p = 0.2265). N = 9-30 per genotype. (L) ASH1L mutant mice have no significant difference in time spent grooming. Data were analyzed by Mann-Whitney test for non-parametric data (p = 0.6315). N = 19-20 per genotype. (M) ASH1L mutant mice have no significant difference in marble burying behavior. Data analyzed by unpaired t-test (t_37_ = 0.8455, p = 0.4032). N = 19-20 per genotype. Representative cages are shown below after marble burying test from WT (left) and *Ash1L^gt/+^* (right) mice.

**Figure S2:**
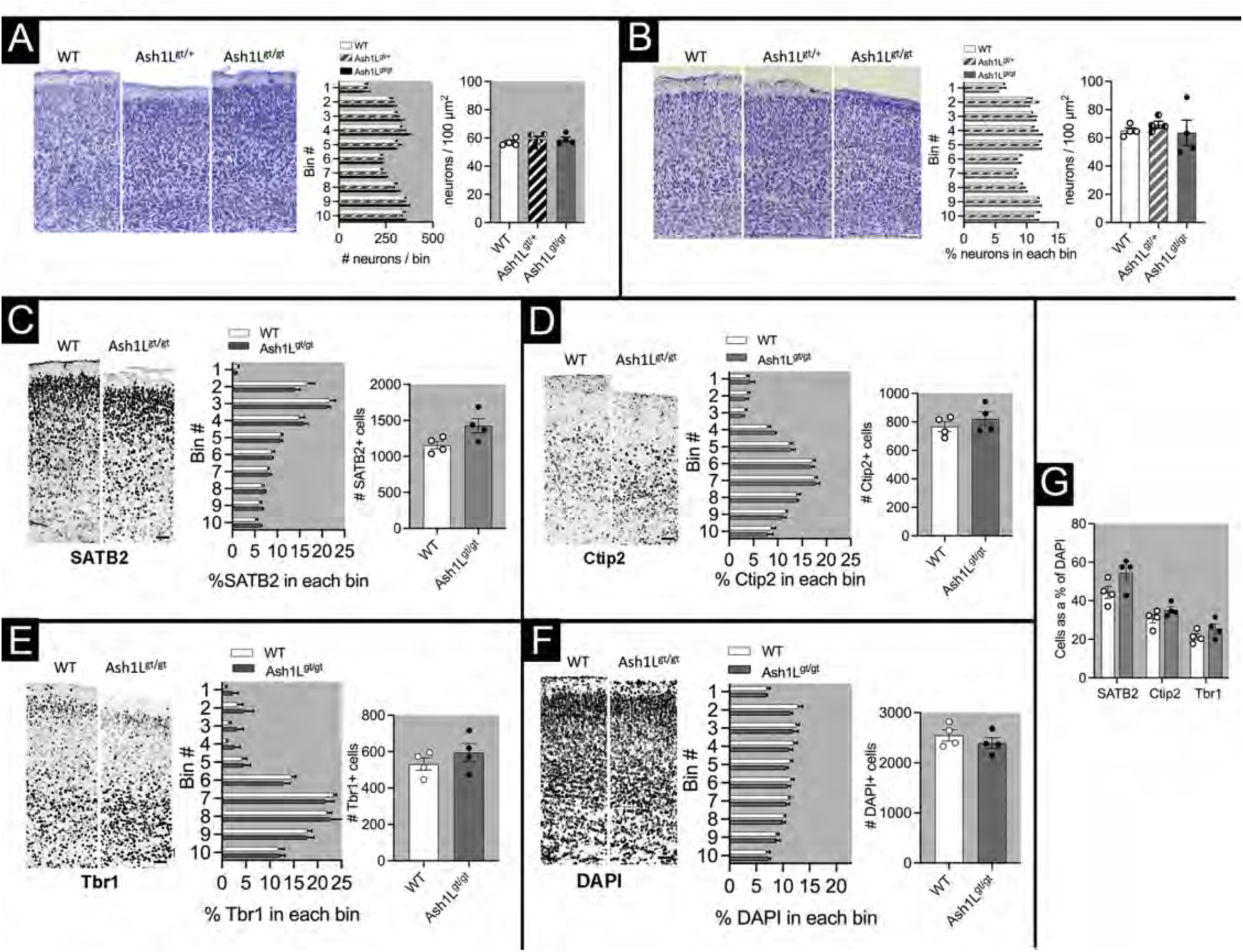
cortical counts in layers and layer markers. (A) Representative images of Nissl-stained cortex at P14 on a B6 background. Scale bar = 100 µm. There is no difference in the distribution of neurons in ASH1L mutant mice on a B6 background. Two-way ANOVA (bin, genotype) finds no effect of genotype (F_2,90_ = 8.998e^-9^, p > 0.9999) or interaction between bin and genotype (F_18,90_ = 0.9835, p = 0.4856). On a B6 background, there is no significant difference in neuron density in ASH1L mutant mice by Kruskal-Wallis test (p=0.6478). (B) Representative images of Nissl-stained cortex at P14 on a CD-1 background. Scale bar = 100 µm. There is no difference in the distribution of neurons in ASH1L mutant mice on a CD-1 background. Two-way ANOVA (bin, genotype) finds no effect of genotype (F_2,108_ = 1.867e^-5^, p > 0.9999) or interaction between bin and genotype (F_18,90_ = 0.6879, p = 0.4862). On a CD-1 background, there is no significant difference in neuron density in ASH1L mutant mice by Kruskal-Wallis test (p=0.4360). N = 4 per genotype (2 males, 2 females). (C) WT and *Ash1L^gt/gt^* cortex at postnatal day 0 is stained for upper-layer neuronal marker SATB2. Scale bar = 50 µm. There is no significant difference in total number (unpaired t-test; t_6_ = 2.389, p = 0. 0.0541) or distribution of SATB2+ neurons in the somatosensory cortex at P0. For neuron distribution, two-way ANOVA (bin, genotype) finds no effect of genotype (F_1,60_ = 2.087e^-18^, p > 0.9999) or interaction between bin and genotype (F_9,60_ = 1.341, p = 0.2354). (D) WT and *Ash1L^gt/gt^* cortex at postnatal day 0 is stained for the lower-layer neuronal marker Ctip2. Scale bar = 50 µm. There is no significant difference in total number (unpaired t-test; t_6_ = 0.9144, p = 0.3958) or distribution of Ctip2+ neurons in the somatosensory cortex at P0. For neuron distribution, two-way ANOVA (bin, genotype) finds no effect of genotype (F_1,60_ = 0.5073, p = 0.4791) or interaction between bin and genotype (F_9,60_ = 0.5235, p = 0.8518). (E) WT and *Ash1L^gt/gt^* cortex at postnatal day 0 is stained for the lower-layer neuronal marker Tbr1. Scale bar = 50 µm. There is no significant difference in total number (unpaired t-test; t_6_ = 1.022, p = 0.3464) or distribution of Tbr1+ neurons in the somatosensory cortex at P0. For neuron distribution, two-way ANOVA (bin, genotype) finds no effect of genotype (F_1,60_ = 7.99e^-18^, p > 0.9999) or interaction between bin and genotype (F_9,60_ = 0.6949, p = 0.7107). (F) WT and *Ash1L^gt/gt^* cortex at postnatal day 0 is stained for the nuclear marker DAPI. Scale bar = 50 µm. There is no significant difference in total number (unpaired t-test; t_6_ = 0.9850, p = 0.3627) or distribution (C) of DAPI+ nuclei in the somatosensory cortex at P0. For nuclear distribution, two-way ANOVA (bin, genotype) finds no effect of genotype (F_1,60_ = 3.710, p = 0.0588) or interaction between bin and genotype (F_9,60_ = 0.4917, p = 0.8745). (G) Number of cells positive for SATB2, Ctip2, and Tbr1 are expressed as a percent of DAPI+ nuclei. By multiple unpaired t-tests, there is no significant difference in the relative percentage of SATB2+ cells (t_6_ = 1.989, p = 0.0939), Ctip2+ cells (t_6_ = 1.610, p = 0.1586), or Tbr1+ cells (t_6_ = 1.465, p = 0.1931).

**Fig. S3:**
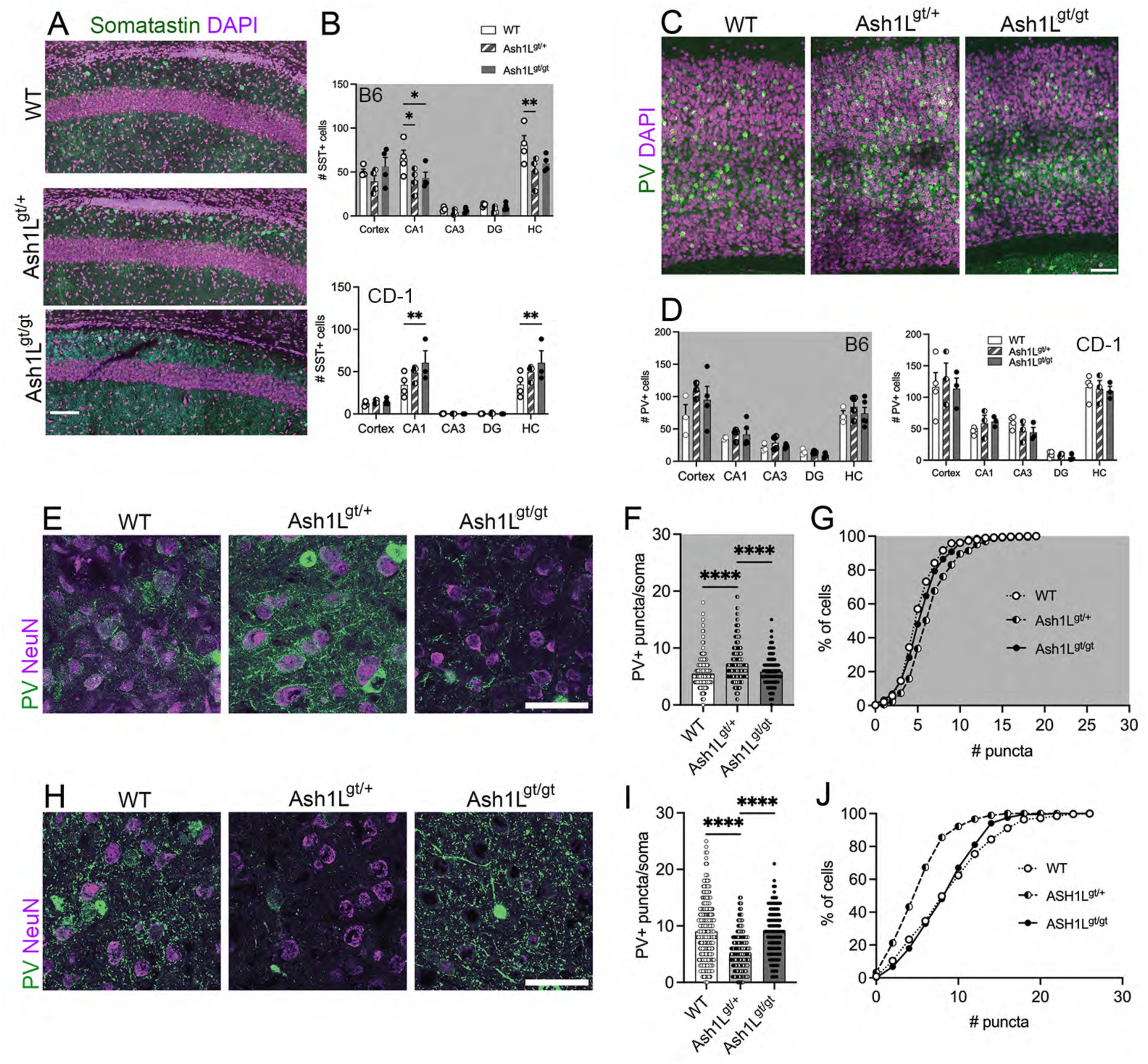
ASH1L mutant mice have changes in inhibitory interneurons in the cortex and hippocampus. (A) Representative images of hippocampus CA1 in CD-1 WT, *Ash1L^gt/+^* and *Ash1L^gt/gt^* mice at postnatal day 14 stained for somatostatin (green) and DAPI (magenta). Scale bar = 100 µm. (B) On a B6 background, *Ash1L^gt/^*^+^ and *Ash1L^gt/gt^* mutant mice have a statistically significant decrease in SST+ cells in CA1 of the hippocampus. p = 0.0017 by two-way ANOVA (brain area, genotype). On a CD-1 background, *Ash1L^gt/+^* and *Ash1L^gt/gt^* mutant mice have a statistically significant increase in SST+ cells in CA1 of the hippocampus. p = 0.009 by two-way ANOVA (brain area, genotype). N = 4 mice per genotype (2 males, 2 females). (C) Representative images of cortex in WT, *Ash1L^gt/+^* and *Ash1L^gt/gt^* mice at postnatal day 14 stained for parvalbumin (PV, green) and DAPI (magenta). Scale bar = 100 µm. (D) On a BL6 background, ASH1L mutant mice have no statistically significant difference in the number of PV+ cells in somatosensory cortex and hippocampal CA1, CA3, and dentate gyrus (DG). On a CD-1 background, ASH1L mutant mice have no statistically significant difference in the number of PV+ cells in somatosensory cortex or hippocampus. (E, H) Representative images of Layer V cortex on B6 (E) and CD-1 (H) backgrounds stained for PV (green) and NeuN (magenta). Scale bar = 50 µm. (F-G) On a B6 background, *Ash1L^gt/+^* neurons show an increased number of perisomatic synapses by PV+ cells onto NeuN+ somas, as compared to WT and *Ash1L^gt/gt^* mice. **** p < 0.0001 by Kruskal-Wallis test with Dunn’s multiple comparisons. (I-J) On a CD-1 background, *Ash1L^gt/+^* neurons show a decreased number of perisomatic synapses by PV+ cells onto NeuN+ somas, as compared to WT and *Ash1L^gt/gt^* mice. **** p <0.0001 by Kruskal-Wallis test with Dunn’s multiple comparisons. N = 96 cells per mouse, 4 mice per genotype (2 males, 2 females).

**Figure S4:** Table 1: Characteristics of human subjects with mutations in ASH1L: We include a table with detailed genetic and phenotypic analysis of subjects in our study including characteristics of the subjects. On all of our subjects, we include genetics (green) neuro-psychiatric symptoms (blue) epilepsy (purple) neurological exam (white), neuro-imaging (gray), medical issues including gastrointestinal symptoms (peach).

**Figure S5:**
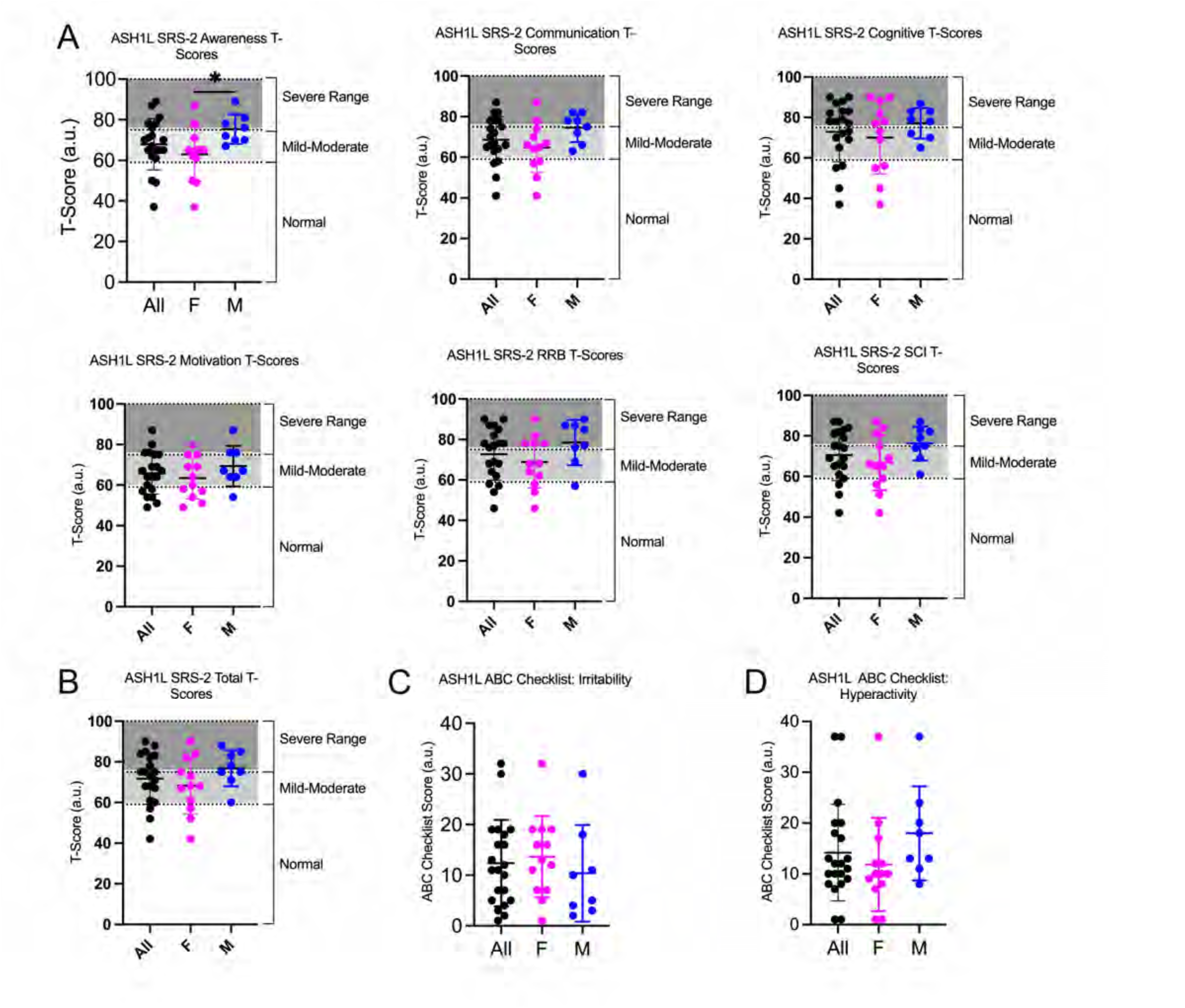
Parental Scales: Social Responsiveness Scale and Aberrant Behavior Scales. A) SRS-2 is a questionnaire filled out by parents used to assess the severity social deficits and symptoms related to autism spectrum disorder (ASD). Mean values of our cohort (n=20) were in mild-moderate range for girls across all domains. However, boys were statistically more severely impacted in terms of social awareness (unpaired t-test, p=0.026) while communication, cognitive, motivation, restricted interests and repetitive behavior (RRB), and social communication and interaction (SCI) scores were not significantly different between girls and boys. (B) The composite scores for SRS-2 is shown and is not significantly different between boys and girls. Aberrant Behavior Checklist (ABC) is a standardized questionnaire that includes an irritability subscale (D) and hyperactivity (E) to measure challenging behaviors. Boys and girls with ASH1L were not significantly different.

**Figure S6:**
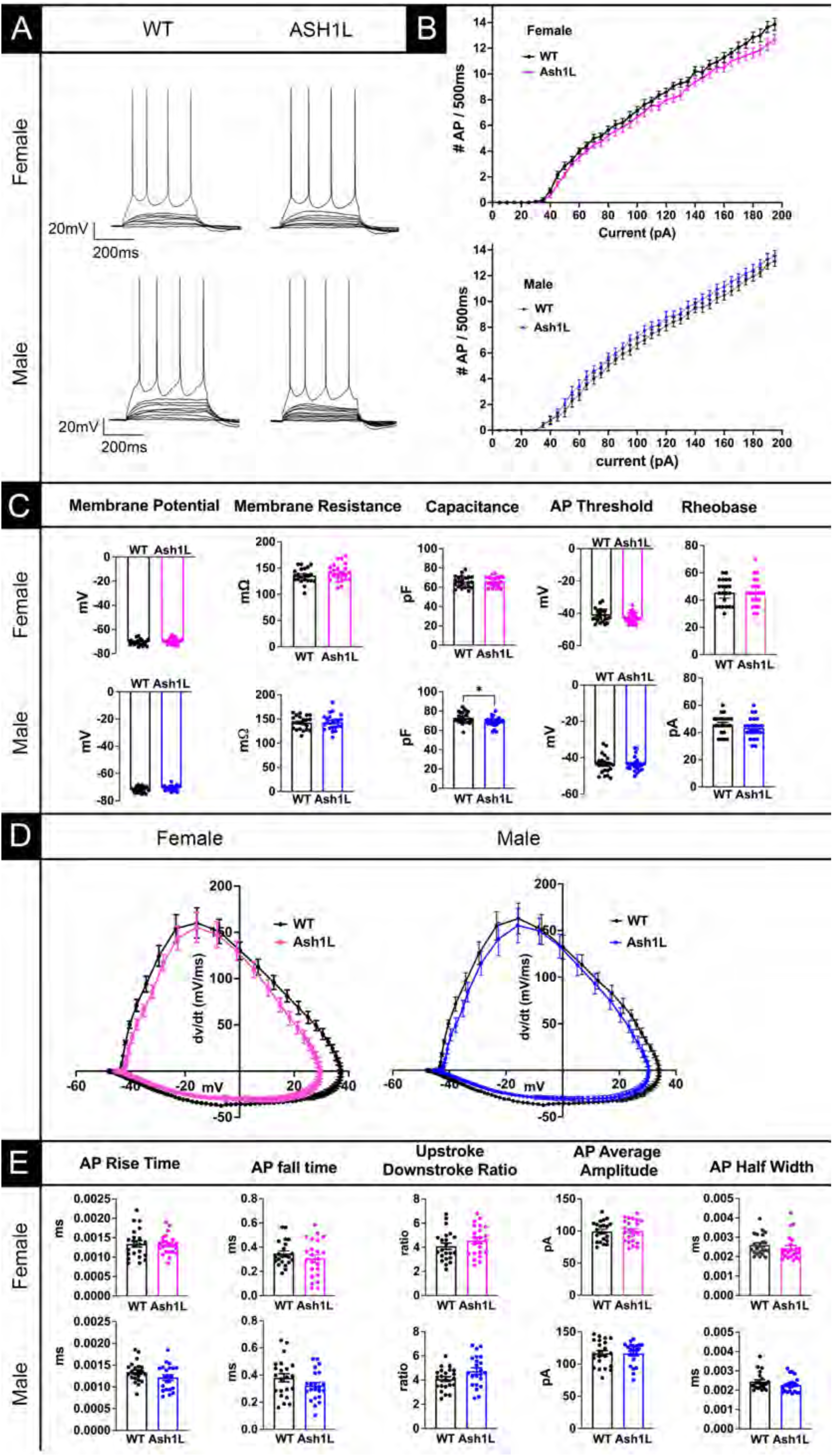
No hyperexcitability in Action Potential Firing in Cortical L5 Pyramidal Neurons from Female Mice. (A) Action Potentials (AP) are triggered by stepwise current injections ranging from -20 to 195 in 5pA steps. (Top) Neurons from *Ash1L^gt/+^* female and male mice fire action potentials at similar amounts of current compared with WT female neurons. (B) Input/ Output curves show no clear separation between female WT and *Ash1L^gt/+^* neuronal AP firing in response to a range of current injection. (C) Intrinsic properties are shown for female and male wild type and *Ash1L^gt/+^* mice: membrane potential, membrane resistance, capacitance, AP threshold potential, and Rheobase (minimum current injection for first AP). (D) Phase plot analysis of AP in wildtype and *Ash1L^gt/+^* female and male mice show few differences in the rate of membrane potential depolarization in female neurons. (E) Analysis of AP properties in cortical neurons from female and mice: AP rise time, fall time, ratio of upstroke to downstroke, amplitude, and half width. Mice were between post-natal day 23-26. n= 8 WT female mice and 22 cells. n= 7 WT male mice and 22 cells. n= 9 for *Ash1L^gt/+^* female mice, and 23 cells. n= 8 *Ash1L^gt/+^* male mice and 22 cells. Data are represented as mean+/-SEM. p values were determined by unpaired t-tests and significant p values less than 0.05 are shown on graphs.

**Figure S7:**
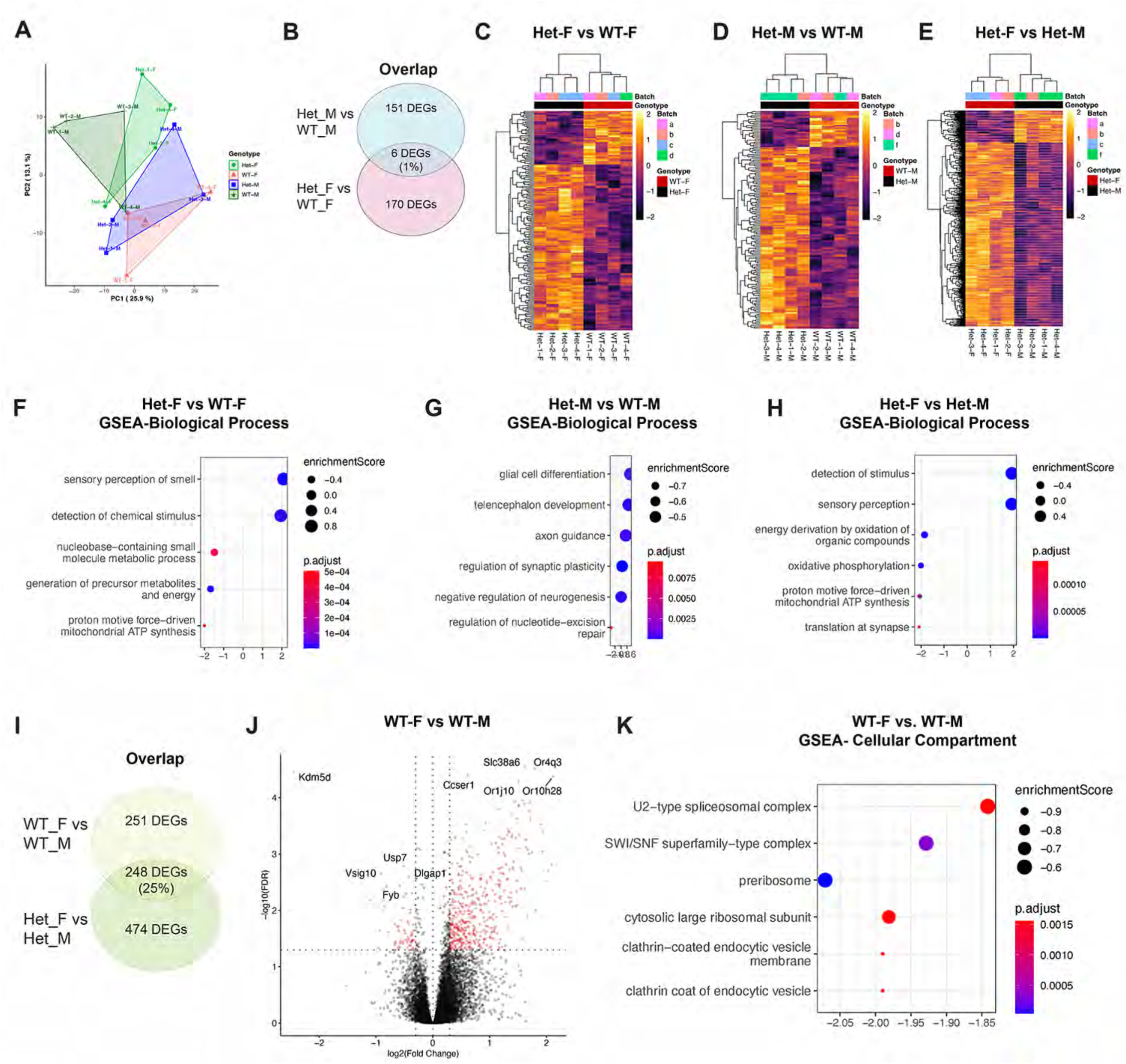
ASH1L modulates sex-specific transcriptional programs. (A) PCA plot shows all different samples in relation to each other. (B) Venn diagrams shows overlap between Het female vs. WT female compared to Het male vs. WT male. (C-E) Heatmaps for the top 100 DEGs for comparisons of: Het female vs. WT female (D), Het male vs. WT male (D) and Het female vs. Het Male (E) show changes in independent biological replicates. Red and black indicate the genotype, and pastel colors indicate individual samples for each genotype. (F-H) Gene set enrichment analysis (GSEA) is shown for biological process for (F) Het female vs. WT female; (G) Het male vs. WT male; and (H) Het female vs. Het Male. Color of circle shows adjusted P value and the size of the circles shows the enrichment scores. (I) Venn diagrams shows overlap between Het female vs. Het Male compared to WT female vs. WT male. (J) Volcano plots showing DEGs above adjusted p value of 0.05 (red dots) that are either downregulated (left side of central dotted lane) or upregulated (right side of central dotted lane). The top 10 DEGs are annotated for WT female vs. WT male. (K) GSEA for cell compartment is shown for WT female vs. WT male. Circle color shows adjusted P value and the size of the circles represents enrichment scores.

**Figure S8:**
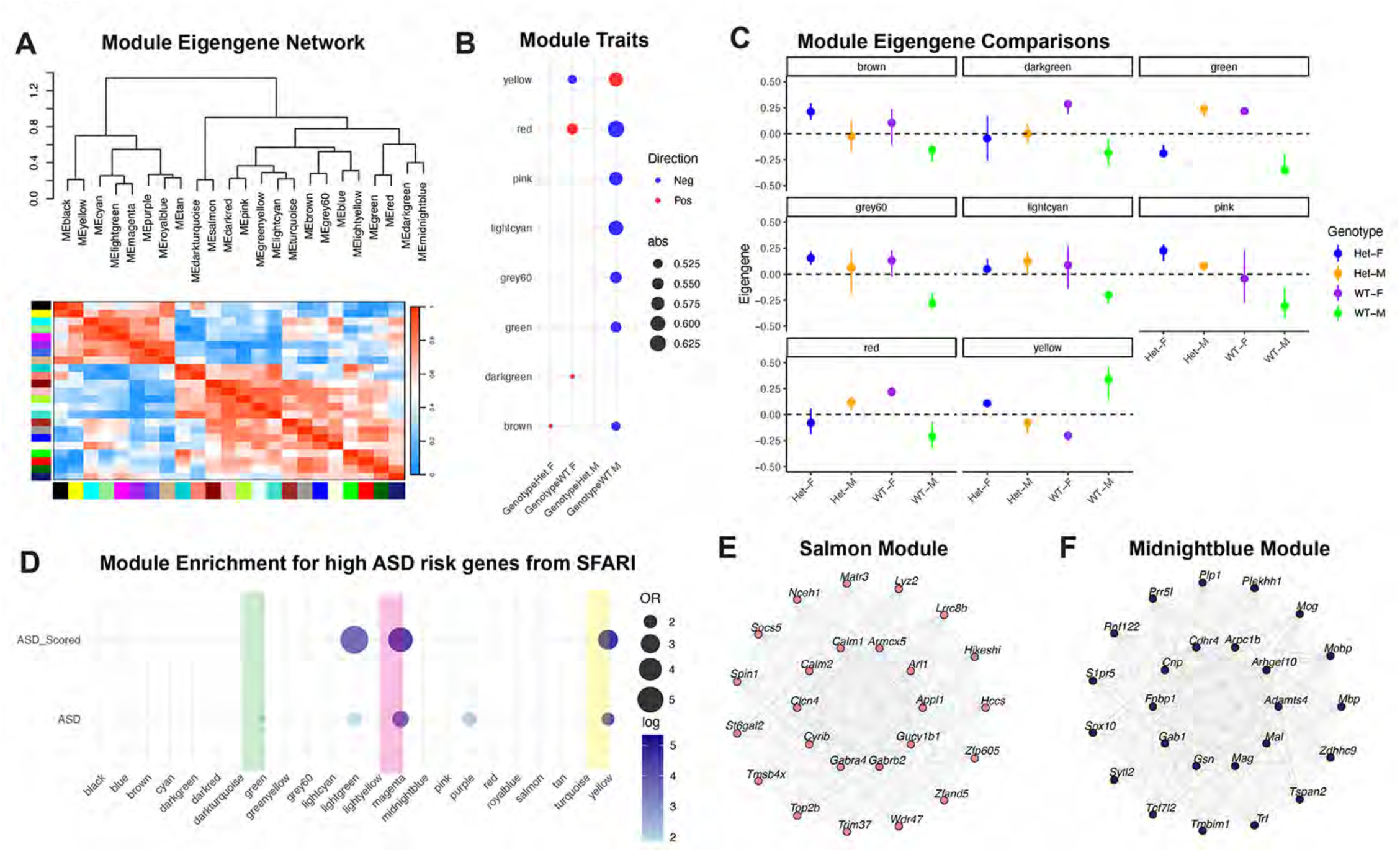
WGCNA shows enrichment for ASD risk genes. (A) Eigengene networks show the relation of every module to each other. (B) WGCNA module trait relationships are shown for HET female, WT female, HET male and WT male. Each module is represented with the direction of gene expression change, indicated as either positive (red) or negative (blue). (C) Eigengene modules are shown for each genotype showing the directionality of the change in expression. (D) WGCNA modules showing enrichment for ASD risk genes based on the SFARI gene database. The size of the solid black circles represents the number of DEGs and the hue color represents the log p-value with darker hues showing greater statistical significance. Solid bars highlight specific modules. (E) Gene networks with relevant hub genes are shown for salmon and (F) midnight blue modules.

